# Stable clique membership in mouse societies requires oxytocin-enabled social sensory states

**DOI:** 10.1101/2025.08.26.672298

**Authors:** Corentin Nelias, Sarah Ghanayem, David Wolf, Marcel Moor, Max F. Scheller, Valery Grinevich, Jonathan R. Reinwald, Wolfgang Kelsch

## Abstract

The ability to form stable de novo relationships in complex environments is essential for social functioning and is impaired in severe psychiatric disorders including autism. Yet, the neurobiological basis and cognitive processes enabling the formation of stable bonds in larger groups remain poorly understood, thereby limiting our ability to develop effective therapies. Here, we establish a semi-naturalistic model of clique formation in mouse societies, where individuals are tracked longitudinally from massive video data. Small, stable rich-clubs develop within these mouse social networks. Consistent with human rich-clubs, these cohesive cliques tended to have high social rank and exerted influence on non-members. Interestingly, neither prior rich-club-membership in a different group nor kinship facilitated entry into rich-clubs. Mimicking sparse population genetics, we probed the open question whether a subtle neuro-cognitive phenotype, namely impaired induction of social sensory processing states by cortical oxytocin signaling, disrupts higher-order social bonding in these complex social environments. Despite preserved social motivation, mice with alterations in this oxytocin subsystem failed to join rich-clubs. They approached group members less consistently, and connections from others towards them fluctuated more as well. This reciprocal disorganization highlights how interactional dynamics within social networks can amplify individual-level deficits, consistent with models of emergent properties of social behavior. These findings underscore the role of oxytocin in tuning sensory systems into a social processing state. Its dysfunction affects an individual’s ability to establish stable relationships in complex social networks, with profound implications for social functioning deficits in psychiatric disorders.

## Introduction

Structured social networks emerge from repeated interactions, leading to preferential associations that may stabilize into enduring affiliative subgroups [1]. Difficulties in establishing stable interpersonal relationships are a hallmark of severe mental disorders and significantly contribute to impairments in everyday functioning [2–5]. The ability to form such de novo social relationships and maintain them also varies substantially across the general population [4, 6, 7]. Importantly, such variations in higher-order social behavior may not necessarily manifest as reduced social interest or interaction frequency [4, 5]. Instead, they often reflect a diminished ability to occupy and maintain functional roles within complex social networks [4, 7]. High-functioning autism spectrum disorder (ASD), often conceptualized as an extreme expression along a continuum of social functioning [4, 6], exemplifies the dissociation between motivation and success in building new relationships [4, 8]. Individuals with ASD show impairments in forming stable relationships across various contexts, even though social motivation is often preserved [4, 6, 9, 10]. These challenges become particularly relevant in adulthood, when social environments become more flexible and require de novo integration.

Network theory provides a powerful framework for understanding the structural underpinnings of social cohesion. It highlights the critical role of tightly connected subgroups in maintaining network resilience and integrity [8, 11, 12]. While such structures are well documented in real-world and digital social networks [4, 8, 11–13], the neurobiological mechanisms that support their formation and maintenance remain poorly understood. In graph theory, highly integrated social configurations can be described as rich-clubs (RC), which are densely interconnected nodes within the networks [8, 13]. These RC support robust social functioning and exert significant influence over information flow, social support, and integration [8, 11, 13, 14]. Identifying how individuals gain stable access to these high-value social structures may provide insight into the subtle cognitive and neural mechanisms regulating social integration.

Social networks are non-linear dynamical systems [8, 13]. This has several implications. Certain phenotypes may be amplified through emergent properties of higher-order social interactions involving multiple individuals [8, 15]. Emergent properties are macro-level phenomena that result from micro-level interactions and cannot be reduced to the sum of their parts. Thus, a change in one individual (or ‘node’) may not only alter its own behavior but also modify the responses of other nodes, leading to dynamic feedback effects [8, 13, 14].

In mammals, oxytocin is a key modulator of social behavior [16–20] and has been investigated as a therapeutic agent for disorders characterized by social deficits [21, 22]. However, clinical trials of intranasal oxytocin have yielded inconsistent and often disappointing outcomes in socially complex settings [21, 22]. This discrepancy raises the question of whether our mechanistic understanding of oxytocin function is sufficient [23], particularly in the context of higher-order social behaviors.

Neuropsychological studies have identified impaired social sensory processing and recognition in the auditory, olfactory and visual domains in ASD [24–27]. Oxytocin modulates diverse region-specific brain functions, including social sensory processing and motivation [16–19]. Oxytocin sets the olfactory system of rodents into a state optimized for processing social information [18, 20]. Oxytocin receptor (OXTR) activation in the anterior olfactory nucleus (AON) increases the signal-to-noise of sensory input [18]. Mice with reduced OXTR signaling in this sensory cortex (OXTR^ΔAON^) are impaired in social recognition at the neuronal and cognitive level [20]. Similarly, human sensory processing is modulated by genetic polymorphisms influencing OXTR expression [28, 29].

It remains unclear what phenotypic consequences oxytocin’s function in sensory state setting has for higher social functioning in real-life conditions. Several scenarios are conceivable. For example, impaired sensory state setting may broadly affect sensory sampling behavior. Alternatively, reduced sensory state modulation may selectively influence social motivation. It is also possible that these subtle sensory deficits become more relevant with increasing complexity, when individuals must learn structured navigation in social networks de novo. The latter scenario would mean that early sensory alterations can cause a phenotype that is typically associated with higher cognitive functions.

To answer these questions, individuals must be observed over longer periods of time in groups. Groups need to be of sufficient size to allow subnetworks to form, stabilize, or dissolve, and to avoid despotic hierarchies typical of smaller groups [30]. Further, animals should be exposed to groups composed of different individuals to disentangle whether internalized prior experience or the specific group context promote RC membership. Moreover, multiple domains of social behavior should be captured to put social relations into a comprehensive perspective. Finally, inspired by natural patterns of genetic variation within populations [31], genetic manipulations should be limited to a minority of animals to ensure that neurotypical individuals continue to shape the group dynamics.

To meet these requirements, we designed a non-invasive, sensor-rich platform (NoSeMaze) for mice [32], which continuously monitors spontaneous social behavior of identified individuals in a larger group over weeks. The sensor-rich system equipped with video and wireless ID tracking, enables automated monitoring of naturalistic interactions and their network dynamics without experimenter intervention. Leveraging the NoSeMaze, we begin here to dissect the factors and mechanisms underlying the formation and stabilization of RC structures. To test how the reduced oxytocin signaling in the olfactory cortex affects complex social behavior, we interspersed a minority of mice carrying OXTR^ΔAON^ into otherwise neurotypical groups.

## Methods

### Animals

All experiments were conducted in adult male mice (C57BL/6J background, Jackson Laboratory). To probe cortical oxytocin signaling, we applied two manipulations: (1) selective, adult-onset deletion of the oxytocin receptor (OXTR) in the anterior olfactory nucleus (AON; OXTR^ΔAON^) via bilateral rAAV1/2-CBA-Cre injection into OXTR_fl/fl_ mice, with control littermates receiving rAAV1/2-CBA-dTomato; and (2) optogenetic activation of oxytocin-releasing neurons in the paraventricular nucleus (PVN) of OXT-Cre mice via rAAV5-DIO-ChR2(H134R)-mCherry. Surgical procedures and histological confirmations followed Wolf et al. [20] and are described in detail in Supplementary Methods. All procedures compiled with the NIH Guide for the Care and Use of Laboratory Animals and EU directive 2010/63, and were approved by the local authority (Regierungspräsidium Karlsruhe, Referat 35, Germany).

### Dyadic social interaction assay

We first assessed social interaction microstructure using a 5-min free-interaction test with an unfamiliar, adolescent, same-sex C57BL/6 partner. OXTR^ΔAON^ and control mice (n = 6 each) had two sessions. Mice for optogenetic experiments had two stimulation sessions (30 Hz pulse trains every 30 s) and two interleaved sham sessions (light blocked). Videos (30 fps) were manually annotated in BORIS [33] into predefined behavioral states (e.g., nose-to-nose sampling, anogenital sampling, approach, idle). Frequencies, durations, and state transition probabilities were compared using linear mixed-effect models (animal ID as random factor).

### NoSeMaze system

The NoSeMaze is a semi-naturalistic environment for continuous, long-term monitoring of social behavior and reinforcement learning in freely interacting mouse groups without human interference [32]. The NoSeMaze comprises a housing arena (nesting, ad libitum food) and an open-field arena (with video-tracking of social interactions, cf. **Fig. 2a**). The two arenas are connected by tubes with RFID readers at both ends for automated identification of passages. Additionally, the open arena is connected to an olfactory reinforcement learning module providing access to water. This module was developed based on the Autonomouse habitat [34]. The full documentation is available at https://github.com/KelschLAB/NoSeMaze. Four parallel NoSeMaze units each housed 9–10 mice for 3-week rounds. Each group included 2–5 OXTR^ΔAON^ mice (see **Supplementary Table S1**) interspersed among neurotypical controls, mimicking sparse genetic variation. Groups were reshuffled between rounds, with most mice participating in multiple rounds. Younger (16–30 weeks, n = 41, n = 16 OXTR^ΔAON^) and older (55–97 weeks, n = 38, n = 10 OXTR^ΔAON^) animals were tested.

### Reward-seeking and social dominance metrics

In addition to video-based behavior (below), behavioral tracking included automated assessment of reinforcement learning at the water lick port and social dominance using RFID-based monitoring in the tubes (cf. **Fig. 5d,f**, see [32], and Supplementary Methods). Social dominance was quantified using David’s scores from automatically detected ‘tube competitions’ (two mice entering from opposite ends; one forced the other to retreat) and by the proportion of initiated and received chases through the tubes.

### Video tracking and event detection

Overhead infrared cameras continuously recorded the open-field arena 24/7. Prior to each round, non-toxic fur bleaching created unique patterns for individual identification of each mouse in the videos. Videos were available for 17 groups (see **Supplementary Table S2**), resulting in a total of approximately 6,100 hours of continuous video data that were analyzed using DeepLabCut (DLC) [35]. DLC models (ResNet-50 backbone) were trained per group to track each mouse, using manually annotated frames from days 1, 8, and 15. Predictions were filtered by likelihood (>0.8) and temporal consistency. Experimenters were fully blind to the genotype. Due to tracking failures, two mice (group 3) and one full group (group 9) were excluded. Data from week 3 of each round were omitted due to fur regrowth. For details, see Supplementary Methods.

Social events were extracted based on spatial proximity using positional data. An ‘interaction’ was defined as any period when two mice were <10 cm apart for ≥1 s (undirected edge). An ‘approach’ was assigned to the mouse that traveled ≥1.5× of its partner’s distance in the 4 s preceding an interaction (directed edge).

### Social networks and ‘rich-clubs’

By tracking all events occurring between each pair of animals on a daily basis, we obtained social networks of dyadic (pairwise) interactions. Interaction networks were undirected; approach networks were directed. Additional undirected networks captured mean interaction duration. Aggregating the first two weeks, we compared OXTR^ΔAON^ vs. controls for (1) total approaches (initiated/received), (2) total interactions, and (3) total interaction time.

To examine higher-order structures, we aggregated daily interaction graphs over three-day windows and pruned them using a mutual nearest-neighbor approach. Specifically, an edge between two animals was retained only if they were among each other’s top k interaction partners (cf. **Fig. 4**). Note that we tested several *k* values (k = 2-4, see Supplement), which all led to similar results (**Supplementary** Fig. 8). All results shown in this article were obtained using *k* = 3.

In these pruned weighted networks, we tested whether animals formed tightly interconnected subgroups, so called ‘rich-clubs’. A rich-club of degree 𝑘 refers to the subset of nodes 𝑣 with degree 𝑑𝑒𝑔(𝑣) ⩾ 𝑘 that are more interconnected than expected by chance.

To assess this, we computed the weighted rich-club coefficient [36], comparing the observed weighted rich-club coefficient 𝜙(𝑘) to that obtained from 1,000 degree-preserving, weight-shuffled random networks, 𝜙_𝑟𝑎𝑛𝑑_(𝑘) (for details and formula, see Supplementary Methods). A normalized coefficient 𝜙_𝑛𝑜𝑟𝑚_(𝑘) > 1 indicates that high-degree nodes not only connect more but also allocate more weight to their mutual links than expected. We were mainly interested in whether rich-clubs remained stable over time. Rich-club stability was assessed in 3-day aggregated networks. We defined an animal as part of the ‘stable rich-club’ (sRC) if it was included in the rich-club in at least 80% of the three-day networks (i.e. 12 out of 15 days). Analyses focused on interaction networks, though approach and interaction-duration networks yielded highly overlapping sRC composition (see **Supplementary** Fig. 7).

Permutation tests (10,000 iterations) assessed whether sRC composition differed from chance. We randomized within-group IDs to test OXTR^ΔAON^ underrepresentation, litter IDs to test sibling co-membership, and, after group reshuffling, IDs in new groups to test whether prior sRC members re-entered sRCs more often than expected (see Supplementary Methods).

### Graph-theoretical metrics

We quantified network properties to compare OXTR^ΔAON^ mice with control animals outside of the sRC and sRC members. Three metrics were computed: out-strength, mean normalized edge fluctuation (NEF), and burstiness. Out-strength reflects the overall number of approaches initiated by an individual. NEF captures how variable an individual’s connections are across partners over time, thereby providing a finer-grained measure of variability than fluctuations in total activity alone. Burstiness quantifies the temporal irregularity of interactions, distinguishing between regular versus clustered contact patterns [37]. NEF and burstiness were calculated separately for incoming and outgoing edges. Group differences were assessed using permutations tests (n = 10,000) based on group medians, accounting for repeated measures (see Supplementary Methods).

## Results

### OXTR deletion in the AON does not alter brief dyadic social interaction patterns

It is not entirely known if loss of OXT signaling in the AON affects the microstructure of self-paced dyadic same-sex interactions. We therefore first analyzed interaction states and the transitions between them in controlled, video-tracked conditions that allow for high resolution mapping of these behaviors (Fig. 1a-b). We benchmarked mice with selective adult deletion of OXTR in the AON (OXTR^ΔAON^ with bilateral injection of AAV_1/2_-CBA-Cre in adult male OXTR_fl/fl_ mice) and their matched controls (Fig. 1c) against a cohort of mice with optogenetics-boosted global OXT release throughout the forebrain (AAV-mediated conditional ChR2 expression in the paraventricular nucleus of male OXT-Cre mice) and sham stimulation in the same animals (Fig.1d).

**Fig. 1:**
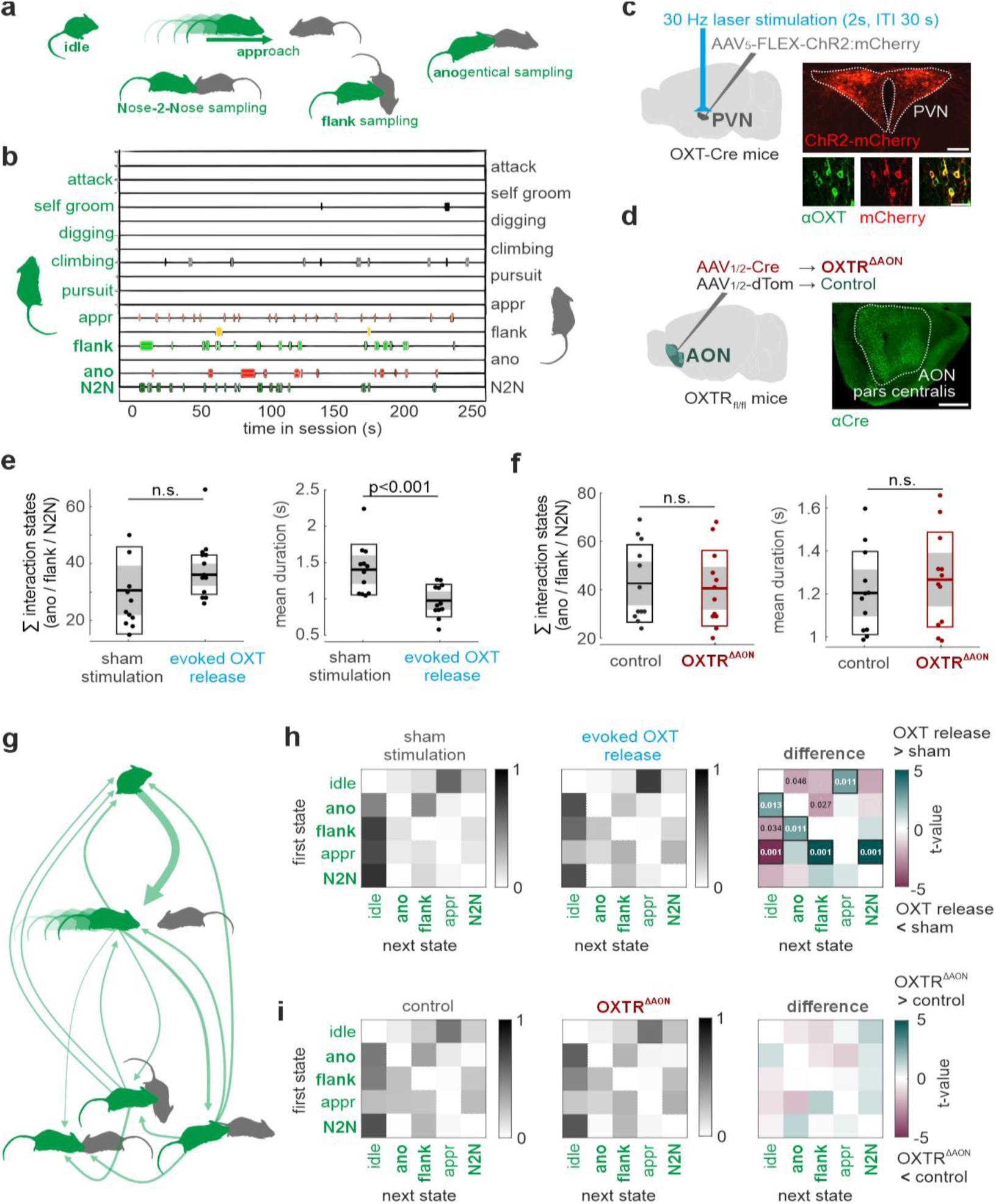
Social microstructure is preserved in OXTR^ΔAON^ mice during brief dyadic interactions. **a)** Ethogram defining key behavioral states observed during self-paced dyadic interactions, including idle (pause), approach, and various forms of social sampling. The experimental mouse (adult) is illustrated in green and the interaction partner in gray. **b)** Example session illustrating the temporal sequence of behavioral states across different interaction categories. **c)** Schematic of optogenetic stimulation and virus injection in the paraventricular nucleus of OXT-Cre mice (left) for conditional expression of ChR2-mCherry in OXT-immunoreactive neurons (right). **d)** Schematic of selective OXTR deletion in the AON of adult OXTR_fl/fl_ mice via AAV_1/2_-Cre injection (left), with Cre expression shown in the AON pars centralis (right). **e)** Even though global optogenetic enhancement of OXT release did not significantly increase the number of sampling events (sum of flank, anogenital, and nose-to-nose; left), it significantly reduced the duration of sampling events (right). **f)** Mice with selective OXTR^ΔAON^ deletion differed neither in the number nor duration of sampling events from matched controls. **g)** Diagram illustrating transitions between behavioral states over time. Arrows indicate the direction and frequency of transitions. **h)** Transition matrices for behavioral state of sham vs. evoked OXT release. Right panel: paired difference matrix (t-values) reveals increased transitions from approach into sampling with boosted OXT release. P-values are FDR-corrected for multiple comparisons. **i)** Transition matrices for control vs. OXTR^ΔAON^ mice show no significant differences in state transitions. Together, these findings indicate that deletion of OXTR in the AON does not disrupt the microstructure or progression of short dyadic social behaviors, supporting the idea that this region contributes more to social sensory processing rather than the initiation of social action patterns. *AON, anterior olfactory nucleus; OXTR, oxytocin receptor; PVN, paraventricular nucleus*

As hypothesized, boosted OXT release led to a tendency toward more sampling events, with significantly shorter durations (Fig. 1e). In contrast, OXTR^ΔAON^ mice did not differ from controls in the number or duration of interaction states (Fig. 1f). Boosted OXT release led to more frequent transitions from approach into sampling behaviors and a reduction of aborted interactions after approaches were initiated (Fig. 1h), consistent with enhanced social engagement. By contrast, OXTR^ΔAON^ mice showed no significant differences from matched controls in transition probabilities (Fig. 1i). Partner behavior remained unchanged across all groups (Supplementary Fig. 1), indicating no reactive changes in partners’ behavior.

In summary, selective deletion of OXTR in the AON did not alter dyadic social interaction behavior during brief interactions, consistent with OXT acting mainly on social cognition in this cortex. We therefore asked what consequences this selective impairment in social cognition may have in complex and dynamic social networks.

### OXTR^ΔAON^ mice display quantitatively normal social activity in naturalistic group settings

To assess social interaction dynamics in a complex naturalistic setting while retaining full tracking of individual behavior, we employed the NoSeMaze system [32]. The NoSeMaze provides a semi-naturalistic sensor-rich environment enabling long-term, automated tracking of cognitive and social behaviors in group-housed mice without human interference (Fig. 2a). We monitored groups of 9–10 RFID-chipped mice (79 total; 22 OXTR^ΔAON^, 57 controls) over continuous three-week rounds in the NoSeMaze. Each group included two to five OXTR^ΔAON^ mice interspersed among controls to mimic sparse genetic variation. The cohort comprised younger (16–30 weeks) and older adults (55–97 weeks) to assure that phenotypes were observed across the adult lifespan (Fig. 2b). To assess which social behaviors depend on specific group configurations, mice lived with different peers in each round of the NoSeMaze (‘reshuffling’, Fig. 2c).

**Fig. 2:**
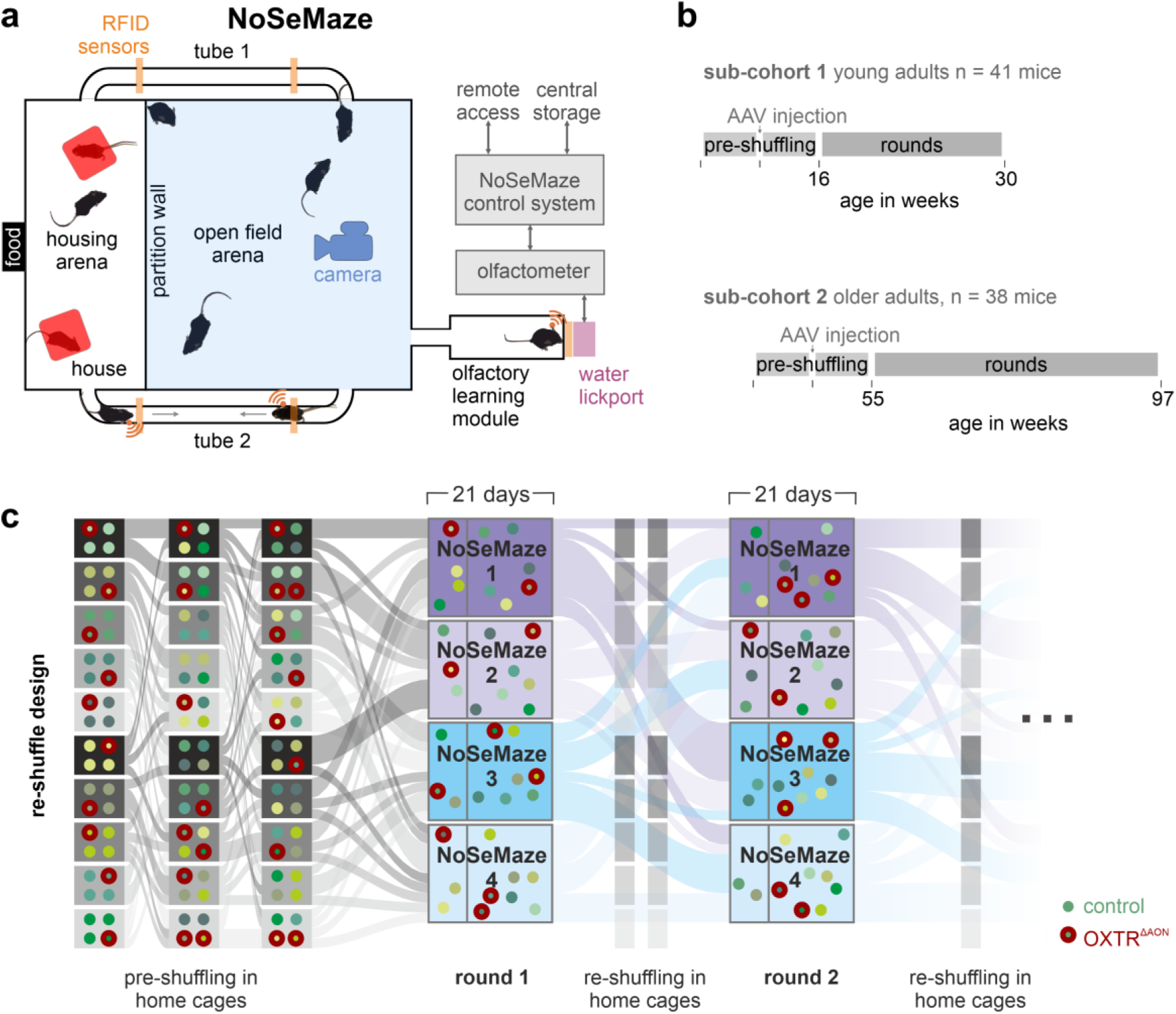
Automated tracking in the NoSeMaze of individuals’ behaviors in groups with interspersed OXTR^ΔAON^ mice. **a)** The NoSeMaze combines ceiling-mounted video surveillance and RFID sensors to continuously track the social behavior, spatial patterns, and reinforcement learning of freely moving mice in complex, group-housed conditions (n = 9–10 mice per group) without human interference. Mice are identified via RFID-tagged implants at access points (e.g., tunnels, water lickport) and tracked by video in the open-field arena to reconstruct social interaction networks from identified individuals. **b)** Experimental timeline for the two sub-cohorts used in this study: younger adults (n = 41; 16 OXTR^ΔAON^, 25 controls) and older adults (n = 38; 10 OXTR^ΔAON^, 28 controls). Following virus injection and pre-shuffling, animals underwent multiple experimental rounds in the NoSeMaze. **c)** Sankey-style diagram illustrating how mice were reassigned across successive NoSeMaze rounds with reshuffling periods in home cages to ensure broad familiarity both before and between rounds. Each group consisted of 9–10 individuals with 2–5 OXTR^ΔAON^ mice. Reshuffling enabled dissociation of effects emerging from internalized traits in individuals versus group composition context. *NoSeMaze, Non-invasive Sensor-rich Maze; RFID, radio-frequency identification*

We then established a pipeline to extract the social interaction networks of the groups of individually identified mice, including interspersed OXTR^ΔAON^ mutants, from continuous video data in the open field arena, across 17 rounds in total (Fig. 3a-b). To identify individual mice, distinct fur patterns were bleached onto the back of the mice before each round (Fig. 3c). We analyzed the first 15 days of each round, as fur regrowth reduced the reliability of pattern recognition by the trained RNN. Before each new round, the fur patterns were refreshed. Approximately 6,100 hours of continuous video data were processed to generate the social networks (Fig. 3d).

**Fig. 3:**
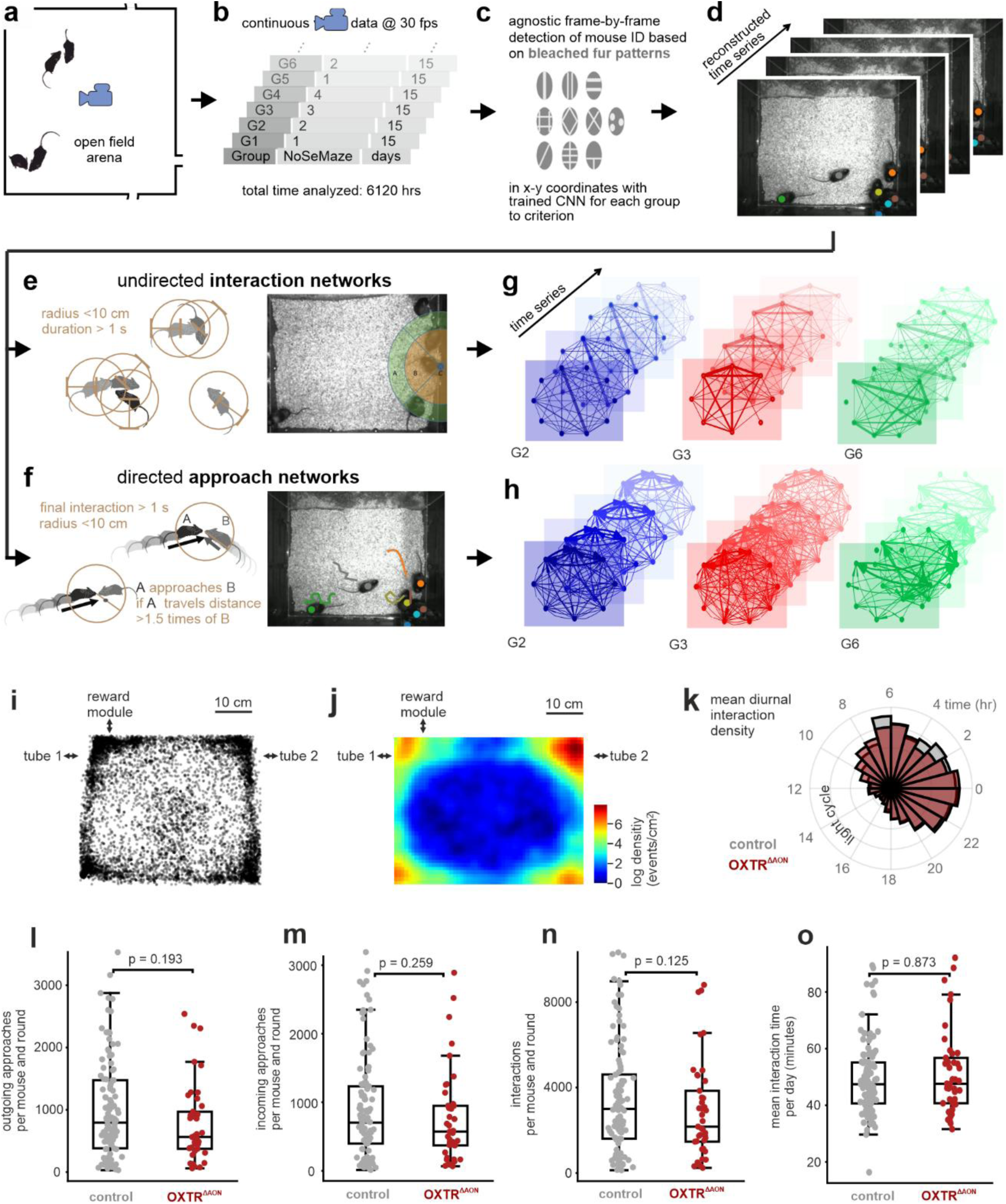
Massive video tracking reveals comparable levels of social interactions in OXTR^ΔAON^ and control mice. **a)** Schematic of the open-field arena in the NoSeMaze, where mice interacted freely. A ceiling-mounted camera continuously recorded video data. **b)** Seventeen groups were recorded over 15 days each, yielding a total of 6,120 hours of video data. **c–d)** Mouse identity and position were extracted frame-by-frame using convolutional neural networks (CNNs) trained on the bleached individual fur patterns on the animals’ backs (see Methods). The CNN output was smoothed to reconstruct individual trajectories over time. **e)** Social interactions were defined as events in which two mice had a distance < 10 cm for at least 1 s. **f)** Directed approaches were identified as a subset of interaction events where one mouse (A) approached another (B) by traveling at least 1.5 times the distance of B in the 4 s preceding their interaction. **g–h)** From these two event types, we constructed time-resolved social networks by aggregating interactions for each 24-h period (see Methods). Interaction networks were undirected, as no distinction was made between the two mice involved **(g)**. In contrast, approach networks were directed, leveraging the direction of approach **(h)**. **i)** Spatial distribution of interaction events in G9 over a round. Each dot represents one interaction. **j)** Heatmap of interaction density (log-transformed), showing clustering around the corners (all NoSeMaze rounds, all mice). **k)** Diurnal distribution of interactions. Mice were most active at night. Overlapping interaction activity was observed for OXTR^ΔAON^ (red) and control (gray) mice. **l–o)** Social behavioral metrics across the first 15 days of each round. Each dot represents a single animal per round. No significant differences were observed between OXTR^ΔAON^ (red) and controls (gray) in **(l)** the number of initiated approaches, **(m)** received approaches, **(n)** total number of interactions, or **(o)** average daily interaction time (in minutes).

The RNN-based pipeline allowed us to extract spatial coordinates for each animal as a function of time. From these tracked positions, we defined and extracted two types of social events: interactions and approaches. This enabled us to represent the data as a social network. Interactions, which were used to build undirected networks, were counted whenever two animals were within less than 10 cm for at least 1.0 s (Fig 3e). Approaches were defined by calculating the distance travelled in the 4s before contact for both interaction partners (Fig. 3f). If the ratio of distance travelled by mouse A relative to mouse B exceeded 1.5, mouse A was considered the approaching individual; otherwise, no approach was counted. Approaches between each mouse pair yielded directed networks. From these measures, we obtained time series of undirected and directed networks (Fig. 3g-h, see Methods).

Spatially, interactions occurred preferentially near the walls (Fig. 3i, Supplementary Fig. 2) and predominantly during the active dark cycle (Fig. 3k). Notably, OXTR^ΔAON^ and control mice displayed quantitatively similar number of interactions and event durations (Fig. 3n-o), as well as comparable numbers of both incoming and outgoing approaches (Fig. 3l-m), suggesting preserved approach behavior in OXTR^ΔAON^ mice. We therefore tested the hypothesis that the social cognition deficits caused by OXTR^ΔAON^ specifically impair the formation of targeted and consistent social relationships.

### Stable rich-clubs emerge in social networks of group-housed mice

Social relationships are dynamic and evolve in time. While they can be unstable or occur by chance, they can also gain stability and persist among some animals that live in the same habitat. We were particularly interested in identifying the key factors that enable the formation of such stable reciprocal cliques within larger groups.

Interaction data were analyzed across five consecutive 3-day windows per round. Given the high baseline connectivity in the networks, graphs were pruned using a mutual nearest-neighbor approach to isolate stable and behaviorally relevant substructures (Fig. 4a; Supplementary Fig. 5 and 6). In most cohorts (81%), we observed the emergence of stable, densely interconnected subnetworks composed of high-degree nodes. These structures (Fig. 4b–c) represent rich-clubs (RC) [14, 18]. This finding was supported by richness coefficients consistently exceeding random chance throughout the experiment (see Supplementary Fig. 4).

RCs that persisted across at least 4 out of 5 three-day-graphs were considered stable rich-clubs (sRC). Notably, sRC were robustly identifiable in both undirected interaction and directed approach networks (Supplementary Fig. 7). Statistical modeling confirmed that such persistence was highly unlikely to occur by chance (p < 10⁻⁷ for pairs; p < 10⁻⁹ for triads), indicating genuine social structure. sRC emerged in all 10 rounds with older adults (Fig. 4c, Supplementary Fig. 5), while in younger adults they were observed in 4 of 7 rounds (Supplementary Fig. 5), potentially suggesting that age may favor clique formation.

sRC members displayed distinct interaction patterns. Compared to non-members, they engaged in more interactions and approaches (Fig. 4e-g), although approach duration did not differ (Fig. 4h). Notably, OXTR^ΔAON^ mice behaved similarly to non-members on these measures (Fig. 4e-h). sRC members also exhibited greater reciprocity: 96.0% of their approaches were reciprocal, compared to 67.7% in the general population (Fig. 4d). Thus, sRCs form in most NoSeMaze groups and can be identified both from undirected interaction graphs and directional approach networks.

**Fig. 4:**
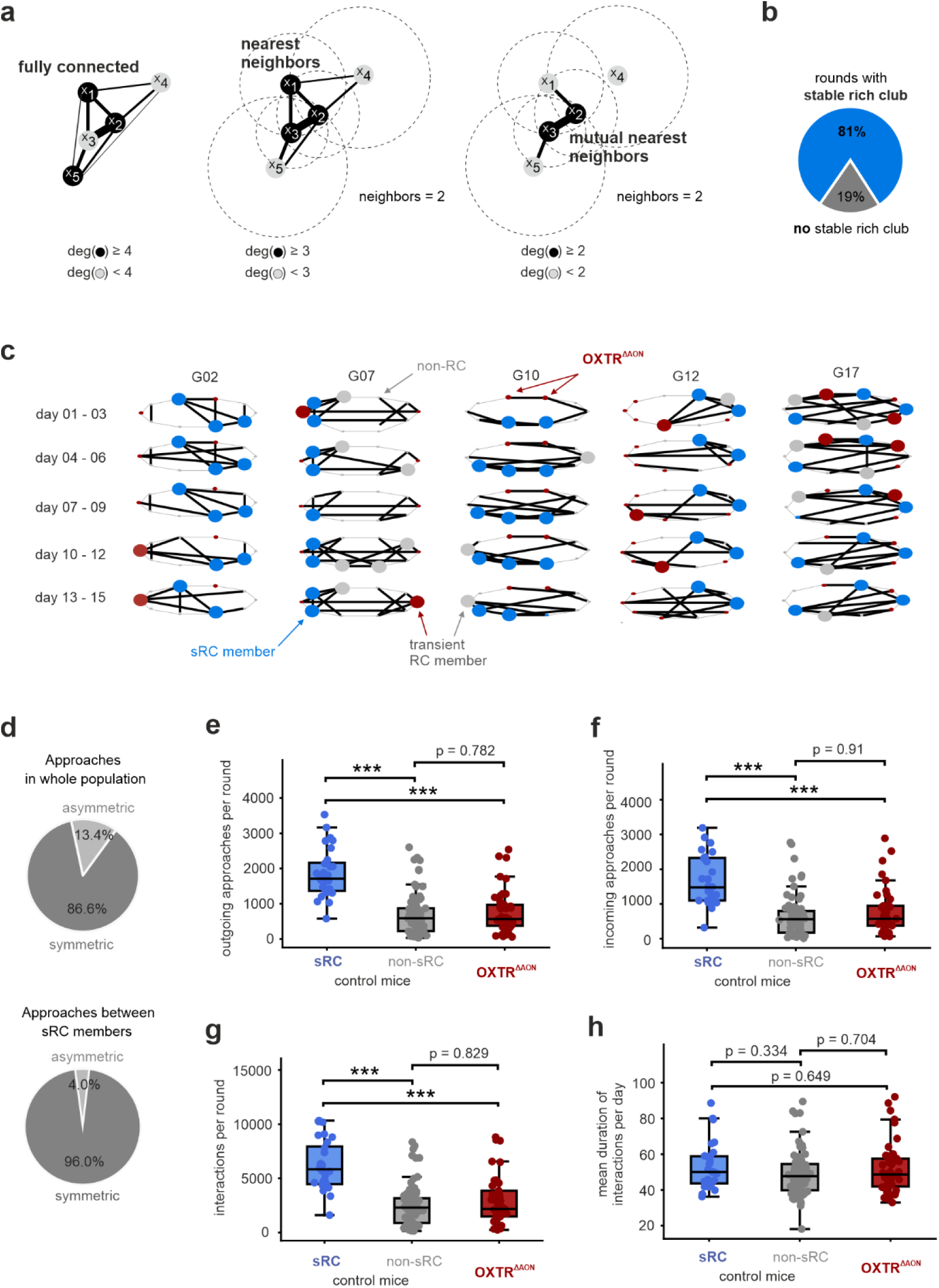
Stable rich clubs (sRCs) emerge in NoSeMaze groups and are characterized by high approach density. **a)** Schematic illustrating the mutual nearest neighbors graph-cut procedure used to identify RCs. Starting from a fully connected graph (left), edges are filtered based on 2^nd^-order nearest neighbors (middle), and further reduced to a final graph (right) that preserves edges only if nodes are mutual 2^nd^-order nearest neighbors of each other. Dashed circles indicate each node’s 2^nd^-nearest neighbor radius; nodes within each other’s circle form mutual connections. Nodes forming RCs are shown in black; others in gray. As the graph-cut criterion becomes more selective, the number of RC members decreases. **b)** Stable RCs were detected in 81% of NoSeMaze rounds (blue). **c)**Representative examples of rich club dynamics across five rounds (G02, G07, G10, G12, and G17) over time. Nodes in ‘stable RCs’ (sRCs), defined as maintaining RC-membership across at least 80% of 3-day time windows, are shown in bold blue; transient members in bold gray. OXTR^ΔAON^ mice are indicated in red. Stability is reflected in the consistent preservation of RC structure over time. **d)** Symmetry of approach behavior of sRC members (bottom) compared to the rest of the population (top). Even more than when examining all mice, sRC members engaged in symmetric approaches among themselves. **e–h)** Quantification of social activity across the three groups: control mice, subdivided into sRC members (blue) and non-sRC members (gray), and OXTR^ΔAON^ mice (red). sRC and non-members only show control mice, ensuring no animal is represented more than once across groups. Each dot represents one animal in one round. sRC members initiated and received significantly more approaches and participated in more interactions than both other groups (***, p < 0.001, permutation test on the median), while OXTR^ΔAON^ mice did not differ from non-member control animals **(e–g)**. Mean daily interaction time did not differ significantly across groups **(h)**.

### Rich-club membership emerges from group-specific dynamics

We next examined potential predictors of sRC membership. Firstly, because sRCs emerge from the interplay among specific individuals, shared upbringing could promote shared sRC membership in adulthood. We took advantage of having multiple siblings per group to test this possibility. Kinship had only a weak, non-significant effect on increasing the likelihood of shared sRC membership beyond random chance (Fig. 5a, Supplementary Fig. 8a).

**Fig. 5:**
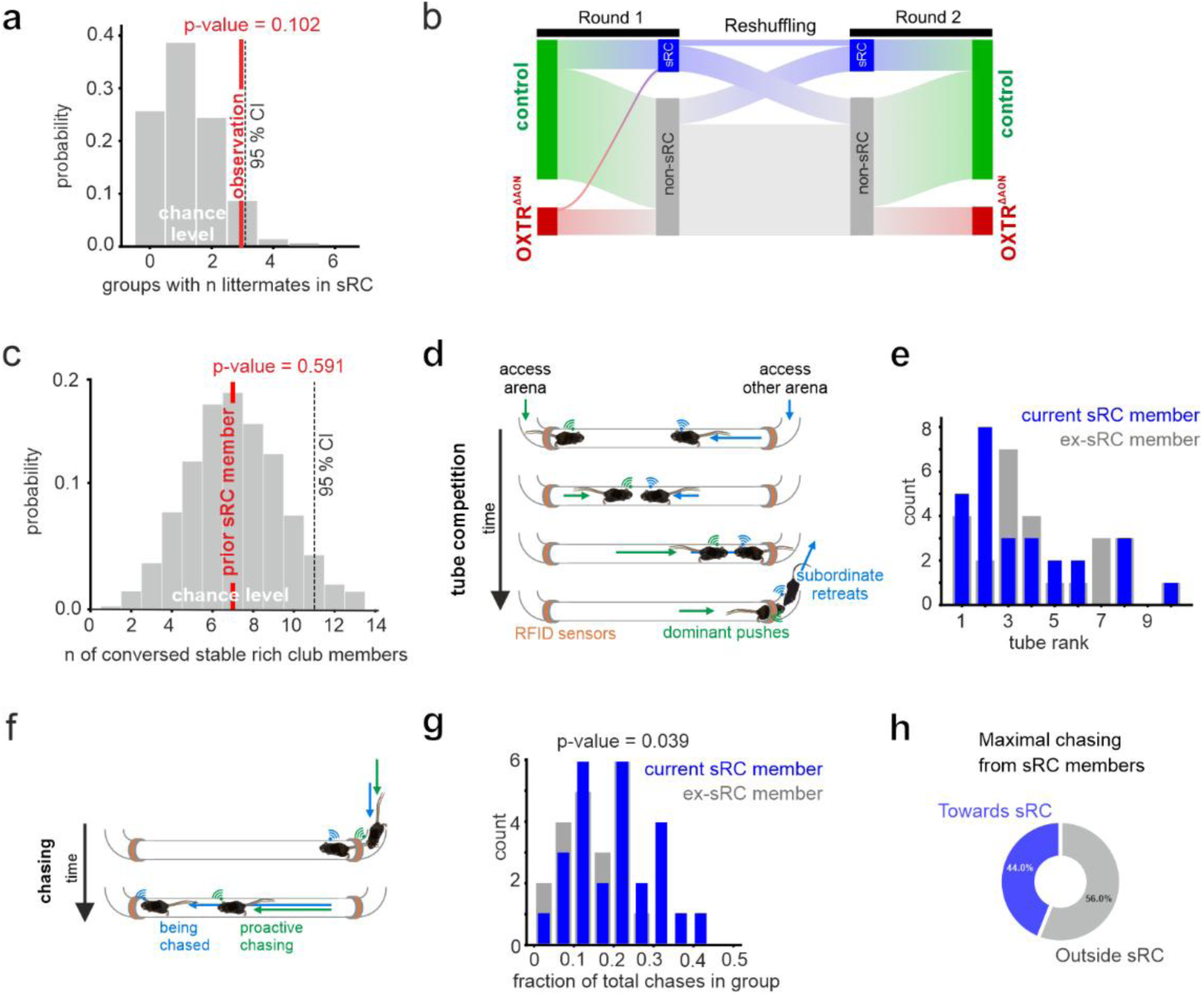
Stable RC membership is shaped by group context and associated with dominance and chasing behavior. **a)** Observed frequency of littermates within the same sRC compared to a null distribution generated by random assignment (see Methods). The observed frequency (red line) falls within the 95% confidence interval of the null model, indicating that shared family background does not significantly influence sRC formation. **b)** Sankey diagram illustrating sRC membership across successive NoSeMaze rounds. The lack of continuity across rounds suggests that sRC membership is not a stable individual trait but group-context dependent. **c)** Probability that former sRC members rejoined an sRC in a different round. The observed number (red line) falls within the null model’s 95% confidence interval, indicating that sRC membership is not an internalized individual characteristic. **d)** Schematic of the tube test used to assess social rank via RFID-tagged outcomes of dyadic competitions. **e)** Tube-test ranks of mice that were sRC members in the current round (blue) and the same individuals in rounds when not in the sRC (ex-sRC members, gray). sRC membership was associated with higher dominance ranks, but was not confined to top-ranking individuals. Likewise, ex-sRC members often retained high ranks even when not in an sRC. High dominance favors, but does not guarantee sRC membership. **f)** Schematic of chasing events detected via the tube system. **g)** Fraction of total chases initiated by current sRC members (blue) and the same individuals when not in the sRC (ex-sRC members, gray). sRC members showed significantly higher chasing activity while in the sRC than when they were not part of the sRC (p = 0.039, permutation test on the mean, 10,000 iterations). High chasing activity may facilitate sRC membership without being a necessary trait. **h)** Directionality of chases performed by sRC members.

Secondly, prior membership experience might promote sRC membership in a subsequent round. Alternatively, sRC may form de novo and be driven by the specific group dynamics. To test this, we leveraged the reshuffling procedure, which generated different group compositions across consecutive NoSeMaze rounds (Fig. 5b). Among the mice participating in multiple NoSeMaze rounds, the probability of re-entering an sRC did not exceed chance (Fig. 5c, Supplementary Fig. 8b). This indicates that sRC membership is not a fixed individual trait, but instead emerges de novo from the specific social dynamics within each group. These findings were robust and held across a range of graph-pruning parameters (see Supplementary, Fig. 8).

Stable RC membership was associated with differences in other social behaviors. sRC members tended to occupy higher ranks in the social hierarchy, which was quantified by incidental tube-test competitions within the NoSeMaze (Fig. 5d-e, Supplementary Fig. 9b). However, the distribution of social ranks varied considerably within sRC members. We also tested for proactive chasing of others through the tubes (Fig. 5f). sRC members initiated significantly more chases than non-members (Fig. 5g, Supplementary Fig. 9e). Notably, sRC members initiated chasing toward both fellow clique members and non-members (Fig. 5h). In contrast to sRC membership (cf. Fig. 5c), social rank and proactive chasing are internalized traits in adult individuals and are largely maintained across different group configurations [32]. Consistently, the correlation of social rank from one NoSeMaze round to the next did not depend on whether the individual was in the sRC or not during that round (Fig. 5b, Supplementary Fig. 10). Together, this suggests that higher ranks facilitate, but are not sufficient for sRC entry.

In summary, most groups in the NoSeMaze formed stable interconnected social structures that can be described as sRCs. We found that mice in the sRCs were significantly more active, engaged in more reciprocal interactions, and initiated more chases than non-members. Because family relationships did not significantly influence rich-club formation, and most mice that entered a club did not rejoin it with new peers, we conclude that sRC formation is an emergent property of the network, reflecting the dynamic nature of social interactions within each group.

### OXTR^ΔAON^ mice fail to integrate into stable rich-clubs

Since the formation of rich-clubs in colonies appears to be a consequence of higher-order social interactions, a crucial question is whether OXTR^ΔAON^ mice enter them or not. OXTR^ΔAON^ mice have been shown to exhibit social recognition deficits [18, 20]. While these deficits did not significantly alter how mice engaged in dyadic interactions (cf. Fig. 3l-o), it is very well possible that OXTR^ΔAON^ mice lack the capacity to engage in complex stable social relationships.

To address this, we compared the number of OXTR^ΔAON^ mice that were able to enter the sRC with the number expected by random chance, using a permutation test. The results (Fig. 6a, Supplementary Fig. 8c) show that OXTR^ΔAON^ mice were significantly less likely to enter the sRC than controls. This impairment was observed in both younger and older adults (see Supplementary Fig. 5).

**Fig. 6:**
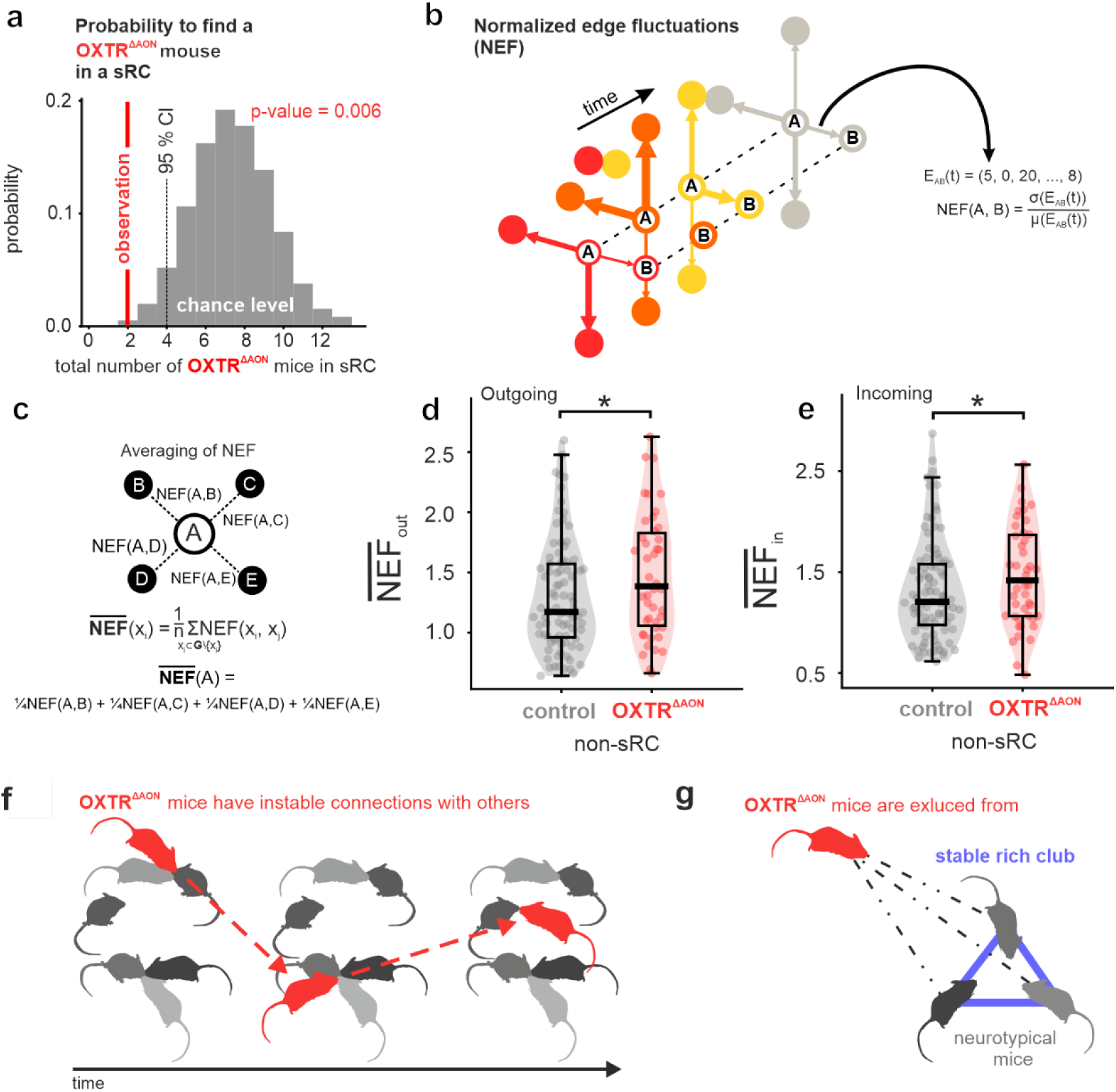
OXTR^ΔAON^ mice fail to integrate into stable rich clubs, display fluctuations in their approach pattern, and receive less consistent input from their peers. **a)** Observed frequency of OXTR^ΔAON^ mice within sRCs compared to a null distribution generated by random assignment (see Methods). The observed frequency (red line) was significantly lower than expected by chance (p=0.006), indicating that OXTR^ΔAON^ mice fail to integrate into sRCs. **b)** Schematic illustrating the normalized edge fluctuations (NEFs), defined for each edge as the coefficient of variation (σ/µ) of its time series. **c)** Illustration of the procedure to obtain node-level averaged NEF by averaging the NEF values of all edges connected to a node. **d–e)** Averaged NEF based on the outgoing (**d**) and incoming (**e**) approaches. OXTR^ΔAON^ mice showed significantly stronger temporal fluctuations in both outgoing and ingoing approaches (p < 0.05, permutation test on the median, 10,000 iterations). **f–g)** Graphical summary: OXTR^ΔAON^ mice exhibit elevated fluctuations in their social approach patterns as well as in the responses from their peers, and are excluded from stable rich clubs. *In all subpanels, each dot represents one animal in one round*.

In summary, OXT signaling in the AON plays a crucial role in the de novo formation of social cliques within larger mouse groups (cf. Fig. 5c). This contrasts with social rank and chasing, which are internalized traits and largely independent of OXT-mediate sensory state setting in adults.

### Unstable network dynamics in OXTR^ΔAON^ mice hinder social integration

Although OXTR^ΔAON^ and control mice did not differ in average measures of interaction time, approach frequency, or social rank (c.f. Fig. 3l-o), OXTR^ΔAON^ mice failed to join sRCs (cf. Fig. 6a). Building on this finding, we asked whether this deficit extends to group-network dynamics, where fluctuating social connections over time might prevent OXTR^ΔAON^ mice from forming and maintaining higher-order social relationships. In other words, even if overall levels of social activity were normal, their ties may have been too inconsistent or erratic to consolidate into stable cliques.

To address this, we examined the temporal fluctuations of directed approach networks. Because current sRC membership itself promotes stability (c.f. Supplementary Fig. 12), we compared OXTR^ΔAON^ mice to control non-sRC-members (who could still be sRC members in other rounds), thereby isolating network features specific to the OXTR^ΔAON^ phenotype. At the global level, the overall strength of outgoing approaches fluctuated to a similar degree in OXTR^ΔAON^ mice and control non-members (Supplementary Fig. 11d), indicating that OXTR^ΔAON^ mice preserved social motivation over time.

To capture fluctuations of the outgoing approaches at the level of specific peers, we required a measure that could handle weighted, fully connected networks rather than reducing them to binary links. Drawing on concepts from the field of temporal graph theory, we developed the mean normalized edge fluctuation (NEF; Fig. 6b-c, see Methods). In simple terms, NEF quantifies how steadily a mouse maintains its connections to others: low NEF indicates consistent ties, while high NEF reflects erratic, highly fluctuating interactions. Formally, for each animal, NEF corresponds to the average coefficient of variation in the temporal weights of its connections to all other mice (Fig. 6c).

Using this measure, we found that OXTR^ΔAON^ mice approached others less consistently than control non-members, as reflected by significantly higher outgoing NEF values (Fig.6d). Thus, although their overall motivation to initiate approaches was preserved, their targeting of specific partners was unstable, leading to locally fluctuating approach patterns over time. Consistently, OXTR^ΔAON^ mice exhibited higher ‘burstiness’ [37] in their approach patterns, with interactions clustering irregularly over time, whereas control non-members approached peers in a steadier, more regular rhythm (Supplementary Fig. 11f).

We next asked whether this instability was also reflected in how OXTR^ΔAON^ mice were approached by others in the group. Indeed, incoming NEF values were higher in OXTR^ΔAON^ mice than in control non-members (Fig. 6e), indicating that peers engaged with them less consistently over time. Incoming burstiness was likewise significantly elevated (Supplementary Fig. 11g), showing that approaches occurred in clustered, irregular bouts rather than steady rhythms.

Together these findings show that OXTR^ΔAON^ mice not only initiate relationships less consistently, but also evoke more variable responses from peers (Fig. 6f). This reciprocal instability prevents their integration into stable rich-clubs (Fig. 6g).

## Discussion

### Summary

This study reveals that mice living in semi-naturalistic social groups form stable and reciprocally connected cliques, so-called ‘stable rich-clubs’ (sRC). These tightly connected hubs reflect genuine and selective affiliative bonds. They represent emergent properties of networks of interacting individuals that influence each other. Embedded within groups of predominantly neurotypical mice, mice with a targeted deletion of the oxytocin receptor in the AON (OXTR^ΔAON^) failed to integrate into these sRC despite normal levels of social activity in both dyadic and group contexts. This reveals that state modulation of social sensory processing through oxytocin becomes critical when structured higher-order interactions must be built in complex social situations.

### Stable rich-clubs

Densely connected hubs such as rich-clubs represent a fundamental component of social network organization in both humans and animals [8, 13, 14]. These structures promote resilience, cohesion, and efficient information flow [1, 4, 5, 7, 10–12, 14]. We focused in mice specifically on sRC, defined as cliques preserved over at least 80% of the lifetime of a group composed of the same individuals. This approach excluded transient structures that may form by chance or fail to stabilize due to shifting affiliative focus. In humans, rich-club members often hold positions of influence and exert strong effects on network dynamics [8, 12, 14]. Consistent with this, sRC member mice tended to occupy higher ranks and displayed broad chasing and approach behavior beyond their immediate cliques, supporting similarly central roles within the group.

### sRC features

Social positions can be determined by intra-individual variables that have been internalized from previous social experiences, so that contextual variables such as the specific group composition play a less important role. In the NoSeMaze system, this applies to social rank and the expression of chasing behavior, both of which are predominantly determined by intra-individual variables and are thus robust against a reshuffling of the group composition [32]. In contrast, belonging to the sRC in one round did not significantly increase the probability of being in the sRC of the next group after reshuffling. Likewise, kinship did not significantly increase the likelihood of belonging to the same clique. This suggests that clique formation is less determined by fixed intra-individual traits and instead emerges anew from the specific interactions and participants within each group. This supports that sRC emerge new in each group, necessitating dynamic learning. The difference between internalized social status and the ability to build novel social relationships also parallels human behavior, where status may facilitate but does not guarantee the ability to form stable contacts [8, 11–13].

### Mice carrying OXTR^ΔAON^ and their deficits

Mice carrying OXTR^ΔAON^ provide a first mechanistic insight in the relation between higher-order social functioning, social processing states and the specific functions of the OXT system. The activation of OXTRs in the AON puts the sensory system into a state of social processing and increases the signal-to-noise ratio in the olfactory bulb [18], which then disseminates information throughout the brain. The consequences of impaired state induction in mice carrying OXTR^ΔAON^ were surprisingly selective in a complex social environment. They kept normal rates of social interactions. Also, the detailed patterns of olfactory social sampling behavior were normal in OXTR^ΔAON^ mice; suggesting together that motivation to interact or sample others are mediated by other OXT-modulated brain circuits. However, mice carrying OXTR^ΔAON^ were markedly impaired in forming stable connections in complex social environments. One might have expected that early sensory impairments caused by OXTR^ΔAON^ would predominantly influence numerous aspects of social behaviors such as motivation, while higher social functions such as mastering stable connections in social networks are a process primarily regulated by higher brain regions. Instead, our findings support a non-hierarchical view of social brain functions, in which selective deficits in higher-order social functions can arise from alterations in early sensory processing rather than from dysfunction in higher brain regions.

### Fluctuations in the interactions of the mice carrying OXTR^ΔAON^

The fluctuating approach patterns of OXTR^ΔAON^ carriers to other mice in the group result in failure to integrate into stable cliques. While overall social activity was preserved, the normalized edge fluctuations and burstiness revealed more erratic and inconsistent patterns of affiliative targeting compared to both control sRC members and non-members. These instabilities likely emerge from deficits in social memory. The oxytocin-induced sensory processing state enables plasticity during interaction, thereby producing reinforced and better differentiated representations of familiar mice in the olfactory cortices [20]. OXTR^ΔAON^ selectively impairs the learning in the social domain, but does not affect non-social olfactory discrimination learning in standard assays [18] and in the NoSeMaze [32]. Importantly, the OXTR deletion occurred only in adult mice, and thus only after mice had already internalized social status [32]. Consequently, OXTR^ΔAON^ carriers largely maintained their social rank in new groups [32], but failed in joining new sRC that requires de novo learning (this study). The deficit from OXTR^ΔAON^ in joining sRC supports that the social sensory state setting actions of oxytocin become particularly relevant in such dynamic situations. These findings underscore the continued role of oxytocin signaling in maintaining coherent social targeting and the formation of new social bonds.

Importantly, these temporal fluctuations also had network-level consequences: neurotypical peers also interacted less consistently with OXTR^ΔAON^ mice, leading to reduced bidirectional stability. This reciprocal disorganization highlights how interactional dynamic in social networks can amplify individual-level deficits, consistent with models of emergent properties of social behavior [8, 12, 14, 15, 38].

### Relations to ASD

This phenotype of OXTR^ΔAON^ mice mirrors some of the key social difficulties in high-functioning ASD. ASD represents the extreme end of a continuum in the capacity to form stable social relationships in the general population [6]. Individuals with ASD may engage in social interactions but often struggle to develop deep, reciprocal relationships in society [4, 5]. Many still function well in rigid, structured environments, but encounter difficulties as they grow older when confronted with increasingly complex and flexible social contexts, particularly when forming friendships [4, 5, 39]. These difficulties affect both personal and professional outcomes [4, 5, 7, 39]. Similarly, OXTR^ΔAON^ mice did not show reduced social interest, but were unable to stabilize their affiliative decisions in complex group situations due to their erratic interaction behavior and the resulting perturbed approach from their peers. Our results therefore suggest that the impairment in OXTR^ΔAON^ mice is not only intrinsic but is amplified by the dynamics of social networks, in which inconsistent behavior can lead to social withdrawal by others. The experimental design, in which a minority of cognitively impaired individuals was embedded within predominantly neurotypical groups, mimicked the sparse patterns of genetic variation commonly found in natural populations and was essential for revealing these network-level amplification effects.

## Conclusion

Taken together, the longitudinal tracking of the collective behavior of individuals in the NoSeMaze enables the identification of mechanisms underlying non-linear affiliative dynamics. The fluctuations of mice carrying OXTR^ΔAON^ in targeting others undermine the formation of stable reciprocal relationships, despite preserved overall social activity. This demonstrates, firstly, how emergent properties from the mutual interactions among subjects in a social environment amplify individual deficits in social cognition. Secondly, these findings highlight the role that early sensory processing can have in the cognition of higher-order social relationships. Finally, it identifies a molecular mechanism underlying the formation of social network organization. These insights thereby provide a framework for the neural basis of social integration and its disruption in psychiatric conditions.

## Data and Code Availability

At the time of publishing, the data will be available upon reasonable request from the corresponding author.

## Acknowledgements

We thank Michael Bram, Jan Ringkamp, and Jens Langejürgen for their help in engineering the NoSeMaze, and Luise Staatsmann for her help in video analyses. We acknowledge the use of ChatGPT (OpenAI, GPT-4, accessed 2025) for assistance with language editing, improving readability, and annotating code. All outputs were reviewed, validated, and verified by the authors, who take full responsibility for the content of this manuscript.

The work was funded by BMBF 3R consortium grants ‘NoSeMaze1’ (161L0277A) and ‘NoSeMaze2’ (16LW0333K) to W.K., Leibniz Association program grant ‘Learning resilience’ (K430/2021) to W.K., Boehringer Ingelheim Foundation grant ‘Complex Systems’ to W.K., BMBF CRCNS grant ‘Oxystate’ (01GQ1708) to W.K, DFG CRC 379 Project C03 to W.K., and the DFG Clinician Scientist Program ‘Interfaces and Interventions in Complex Chronic Conditions’ (EB187/8-1) to J.R.

## References

1. Kappeler PM, Clutton-Brock T, Shultz S, Lukas D. Social complexity: patterns, processes, and evolution. Behav Ecol Sociobiol. 2019;73:5.

2. Adolphs R. The social brain: neural basis of social knowledge. Annu Rev Psychol. 2009;60:693–716.

3. Frith CD, Frith U. The neural basis of mentalizing. Neuron. 2006;50:531– 534.

4. Kasari C, Locke J, Gulsrud A, Rotheram-Fuller E. Social networks and friendships at school: comparing children with and without ASD. J Autism Dev Disord. 2011;41:533–544.

5. Howlin P, Goode S, Hutton J, Rutter M. Adult outcome for children with autism. J Child Psychol Psychiatry. 2004;45:212–229.

6. Baron-Cohen S. Theory of mind and autism: A review. Int. Rev. Res. Ment. Retard., vol. 23, Academic Press; 2000. p. 169–184.

7. Holt-Lunstad J, Smith TB, Layton JB. Social Relationships and Mortality Risk: A Meta-analytic Review. PLOS Med. 2010;7:e1000316.

8. Dunbar RI, Spoors M. Social networks, support cliques, and kinship. Hum Nat Hawthorne N. 1995;6:273–290.

9. Chevallier C, Kohls G, Troiani V, Brodkin ES, Schultz RT. The social motivation theory of autism. Trends Cogn Sci. 2012;16:231–239.

10. Pelphrey KA, Carter EJ. Brain mechanisms for social perception: lessons from autism and typical development. Ann N Y Acad Sci. 2008;1145:283– 299.

11. Borondo J, Morales AJ, Benito RM, Losada JC. Multiple leaders on a multilayer social media. Chaos Solitons Fractals. 2015;72:90–98.

12. Centola D. The spread of behavior in an online social network experiment. Science. 2010;329:1194–1197.

13. Colizza V, Flammini A, Serrano MA, Vespignani A. Detecting rich-club ordering in complex networks. Nat Phys. 2006;2:110–115.

14. Dong Y, Tang J, Chawla NV, Lou T, Yang Y, Wang B. Inferring social status and rich club effects in enterprise communication networks. PloS One. 2015;10:e0119446.

15. Johnson CW. What are emergent properties and how do they affect the engineering of complex systems? Reliab Eng Syst Saf. 2006;91:1475–1481.

16. Ferguson JN, Young LJ, Hearn EF, Matzuk MM, Insel TR, Winslow JT. Social amnesia in mice lacking the oxytocin gene. Nat Genet. 2000;25:284– 288.

17. Marlin BJ, Mitre M, D’amour JA, Chao MV, Froemke RC. Oxytocin enables maternal behaviour by balancing cortical inhibition. Nature. 2015;520:499–504.

18. Oettl L-L, Ravi N, Schneider M, Scheller MF, Schneider P, Mitre M, et al. Oxytocin Enhances Social Recognition by Modulating Cortical Control of Early Olfactory Processing. Neuron. 2016;90:609–621.

19. Walum H, Young LJ. The neural mechanisms and circuitry of the pair bond. Nat Rev Neurosci. 2018;19:643–654.

20. Wolf D, Hartig R, Zhuo Y, Scheller MF, Articus M, Moor M, et al. Oxytocin induces the formation of distinctive cortical representations and cognitions biased toward familiar mice. Nat Commun. 2024;15:6274.

21. Guastella AJ, MacLeod C. A critical review of the influence of oxytocin nasal spray on social cognition in humans: evidence and future directions. Horm Behav. 2012;61:410–418.

22. Yamasue H, Okada T, Munesue T, Kuroda M, Fujioka T, Uno Y, et al. Effect of intranasal oxytocin on the core social symptoms of autism spectrum disorder: a randomized clinical trial. Mol Psychiatry. 2020;25:1849–1858.

23. Leng G, Ludwig M. Intranasal Oxytocin: Myths and Delusions. Biol Psychiatry. 2016;79:243–250.

24. Endevelt-Shapira Y, Perl O, Ravia A, Amir D, Eisen A, Bezalel V, et al. Altered responses to social chemosignals in autism spectrum disorder. Nat Neurosci. 2018;21:111–119.

25. Joseph RM, Tanaka J. Holistic and part-based face recognition in children with autism. J Child Psychol Psychiatry. 2003;44:529–542.

26. Kendrick KM, Lévy F, Keverne EB. Changes in the sensory processing of olfactory signals induced by birth in sheep. Science. 1992;256:833–836.

27. Rozenkrantz L, Zachor D, Heller I, Plotkin A, Weissbrod A, Snitz K, et al. A Mechanistic Link between Olfaction and Autism Spectrum Disorder. Curr Biol CB. 2015;25:1904–1910.

28. LoParo D, Waldman ID. The oxytocin receptor gene (OXTR) is associated with autism spectrum disorder: a meta-analysis. Mol Psychiatry. 2015;20:640–646.

29. Skuse DH, Lori A, Cubells JF, Lee I, Conneely KN, Puura K, et al. Common polymorphism in the oxytocin receptor gene (OXTR) is associated with human social recognition skills. Proc Natl Acad Sci U S A. 2014;111:1987–1992.

30. Sofer Y, Zilkha N, Gimpel E, Wagner S, Chuartzman SG, Kimchi T. Sexually dimorphic oxytocin circuits drive intragroup social conflict and aggression in wild house mice. Nat Neurosci. 2024;27:1565–1573.

31. Selander RK. Behavior and Genetic Variation in Natural Populations. Am Zool. 1970;10:53–66.

32. Reinwald, JR, Ghanayem, S, Wolf, D, Lebedeva, J, Lebhardt, P, Gölz. O, et al. Individual differences drive social hierarchies in mouse societies. bioRxiv. 2025. 2025.

33. Friard O, Gamba M. BORIS: a free, versatile open-source event-logging software for video/audio coding and live observations. Methods Ecol Evol. 2016;7:1325–1330.

34. Erskine A, Bus T, Herb JT, Schaefer AT. AutonoMouse: High throughput operant conditioning reveals progressive impairment with graded olfactory bulb lesions. PloS One. 2019;14:e0211571.

35. Mathis A, Mamidanna P, Cury KM, Abe T, Murthy VN, Mathis MW, et al. DeepLabCut: markerless pose estimation of user-defined body parts with deep learning. Nat Neurosci. 2018;21:1281–1289.

36. Alstott J, Panzarasa P, Rubinov M, Bullmore ET, Vértes PE. A unifying framework for measuring weighted rich clubs. Sci Rep. 2014;4:7258.

37. Goh K-I, Barabási A-L. Burstiness and memory in complex systems. Europhys Lett. 2008;81:48002.

38. Freund J, Brandmaier AM, Lewejohann L, Kirste I, Kritzler M, Krüger A, et al. Emergence of individuality in genetically identical mice. Science. 2013;340:756–759.

39. Chan DV, Doran JD, Galobardi OD. Beyond Friendship: The Spectrum of Social Participation of Autistic Adults. J Autism Dev Disord. 2023;53:424–437.

