## Supplementary Material for "Stable clique membership in mouse societies requires oxytocin-enabled social sensory states"

### Supplementary Figures

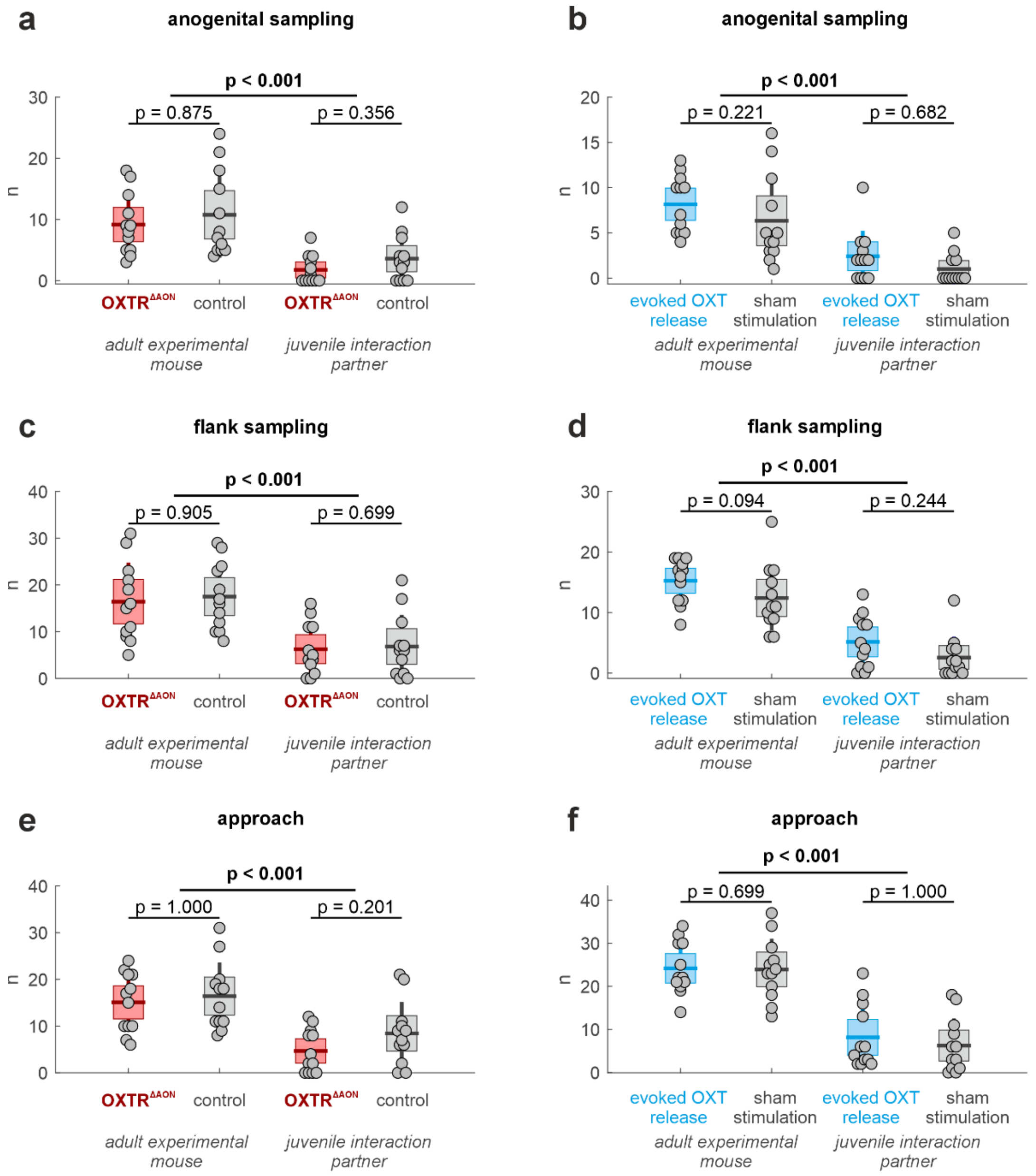


**Supplementary Fig. 1: Sampling and approach behaviors in adult experimental mice and their juvenile partners during self-paced dyadic interactions.**

*OXTR$^{\Delta AON}$ vs. control sessions (left):* Frequency of anogenital sampling (**a**), flank sampling (**c**), and approach behavior (**e**) initiated by adult experimental mice (OXTR$^{\Delta AON}$ vs. control; left panels) and by their respective juvenile interaction partners (when paired with OXTR$^{\Delta AON}$ or control adults; right panels). Behaviors of adult experimental mice and juvenile interaction partners were quantified separately and compared across genotypes of the experimental animal. No significant differences were observed across genotypes in either group (two-sided permutation test on the median, n = 10,000 permutations). However, across all categories, adult mice exhibited higher levels of social sampling than their juvenile partners.

*Evoked OXT release vs. sham stimulation sessions (right):* Frequency of anogenital sampling (**b**), flank sampling (**d**), and approach behavior (**f**) in mice undergoing optogenetically evoked oxytocin (OXT) release compared to sham stimulation. Data are shown separately for adult experimental mice (left panels) and their juvenile interaction partners (right panels).

Each dot represents one individual per session.

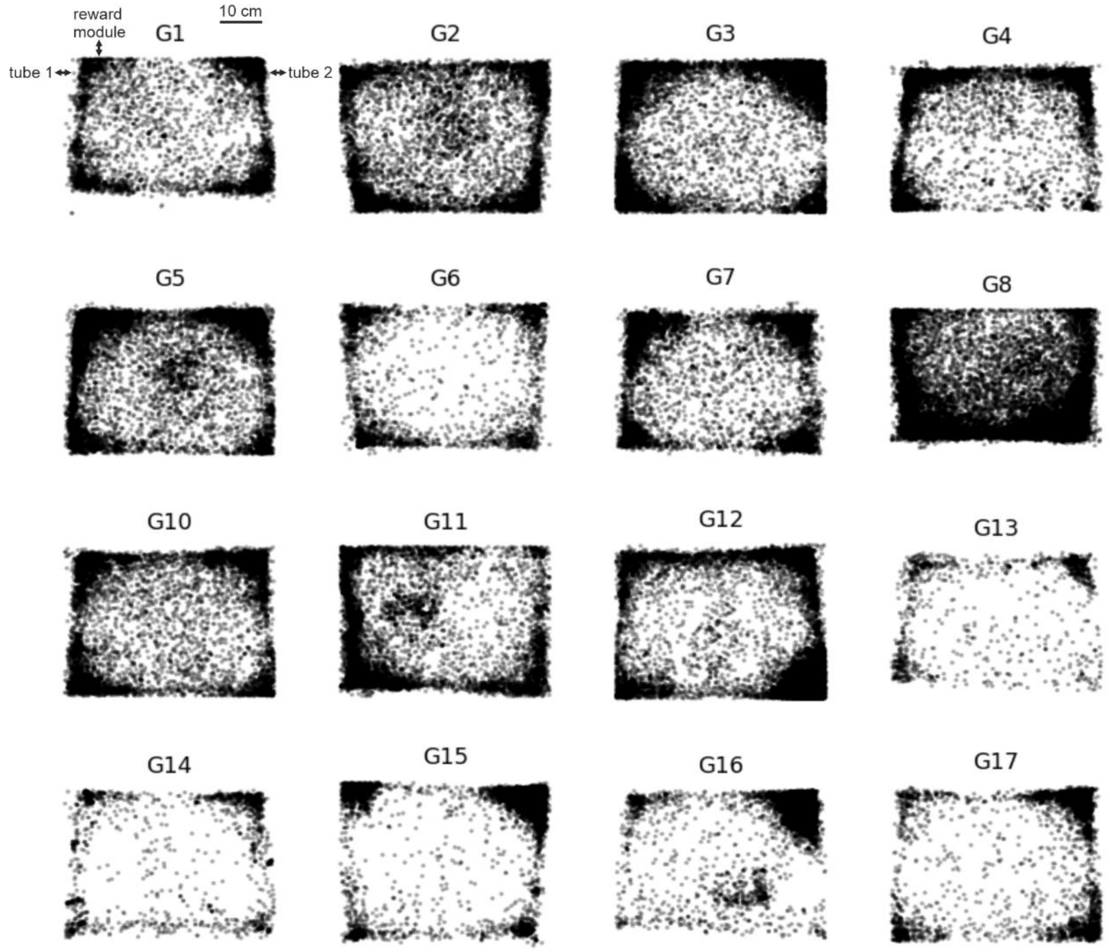


**Supplementary Fig. 2: Spatial distribution of interaction events across all 16 NoSeMaze groups.**
Each panel (G1–G8, G10-G17; G9 excluded, see Methods) shows the spatial locations of individual interaction events detected over one full NoSeMaze round for a given group. Each dot represents a social interaction defined by physical proximity (<10 cm for ≥1 sec) between two individually tracked mice. Interaction activity was most pronounced around the corners and along the arena walls, particularly near access points such as entry tubes and the reward module (top wall).

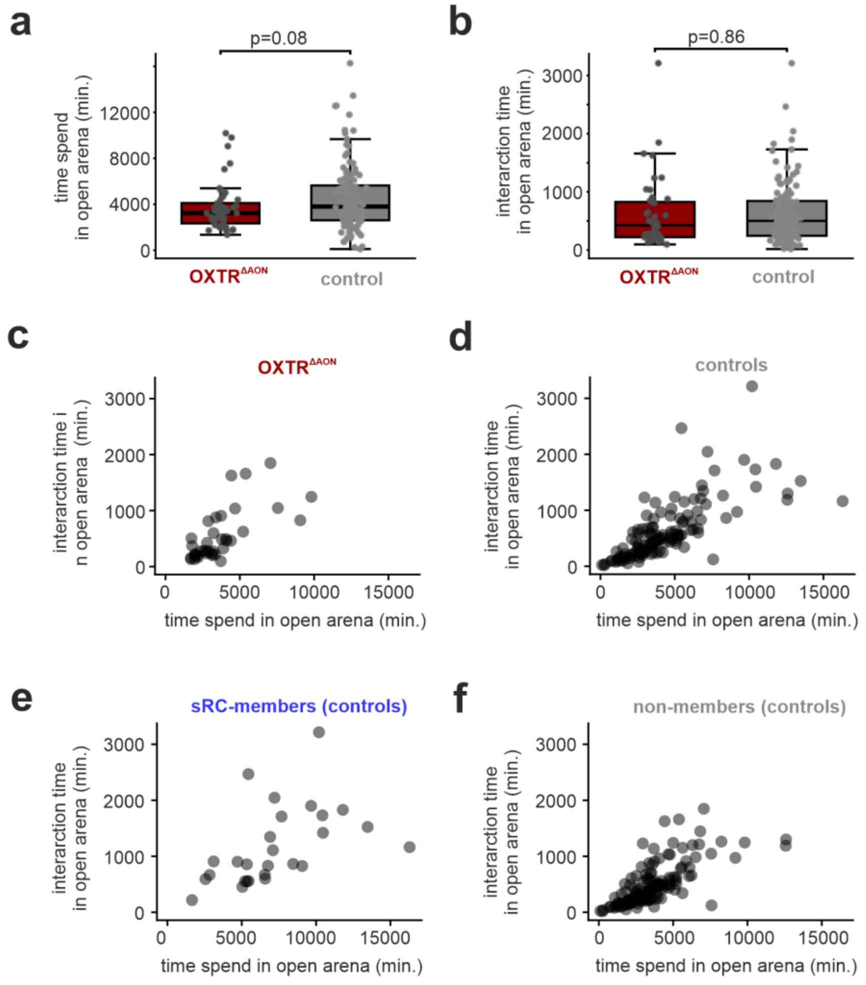


**Supplementary Fig. 3: Time spent in the open arena and social interaction time in OXTR$^{\Delta AON}$ and control mice.**

**a)** Total time spent in the open arena per round was slightly reduced in OXTR$^{\Delta AON}$ compared to control mice ($p = 0.08$, two sided permutation test on the mean with 10,000 permutations accounting for animal identity). **b)** Total time spent in social interactions in the open arena did not differ significantly between OXTR$^{\Delta AON}$ and control mice ($p = 0.86$, two sided permutation test on the mean with 10,000 permutations accounting for animal identity). **c–f)** Correlations between total time spent in the open arena and interaction time was similar across all groups: **c)** OXTR$^{\Delta AON}$ mice; **d)** all control mice; **e)** control sRC-members; **f)** control non-sRC-members.

Each dot represents one individual per session.

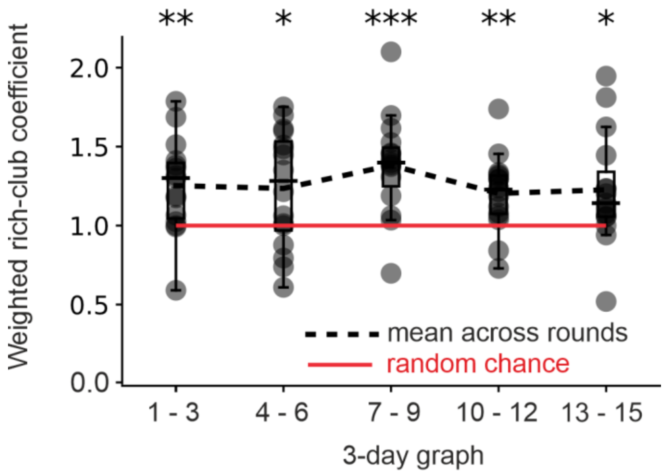


**Supplementary Fig. 4: Sustained presence of weighted rich-club organization across time.**
The normalized weighted rich-club coefficient ($\phi_{norm}(k)$) is shown across five consecutive 3-day windows spanning the 15-day observation period, for nodes of degree k ≥ 3. Each dot represents the normalized weighted rich-club coefficient for one NoSeMaze group; box plots indicate the distribution (median and interquartile range). The dashed black line shows the average rich-club coefficient across groups; the red line marks the null expectation ($\phi_{norm}(k) = 1$) derived from randomized control networks. A normalized rich-club coefficient greater than 1 indicates that high-degree nodes (i.e., animals with many social connections) are more strongly interconnected and allocate disproportionately more weight to their mutual interactions than expected by chance (see methods). Asterisks denote significant deviation from the null expectation (one sample t-test; * : $p < 0.05$, ** : $p < 0.01$, *** : $p < 0.001$).

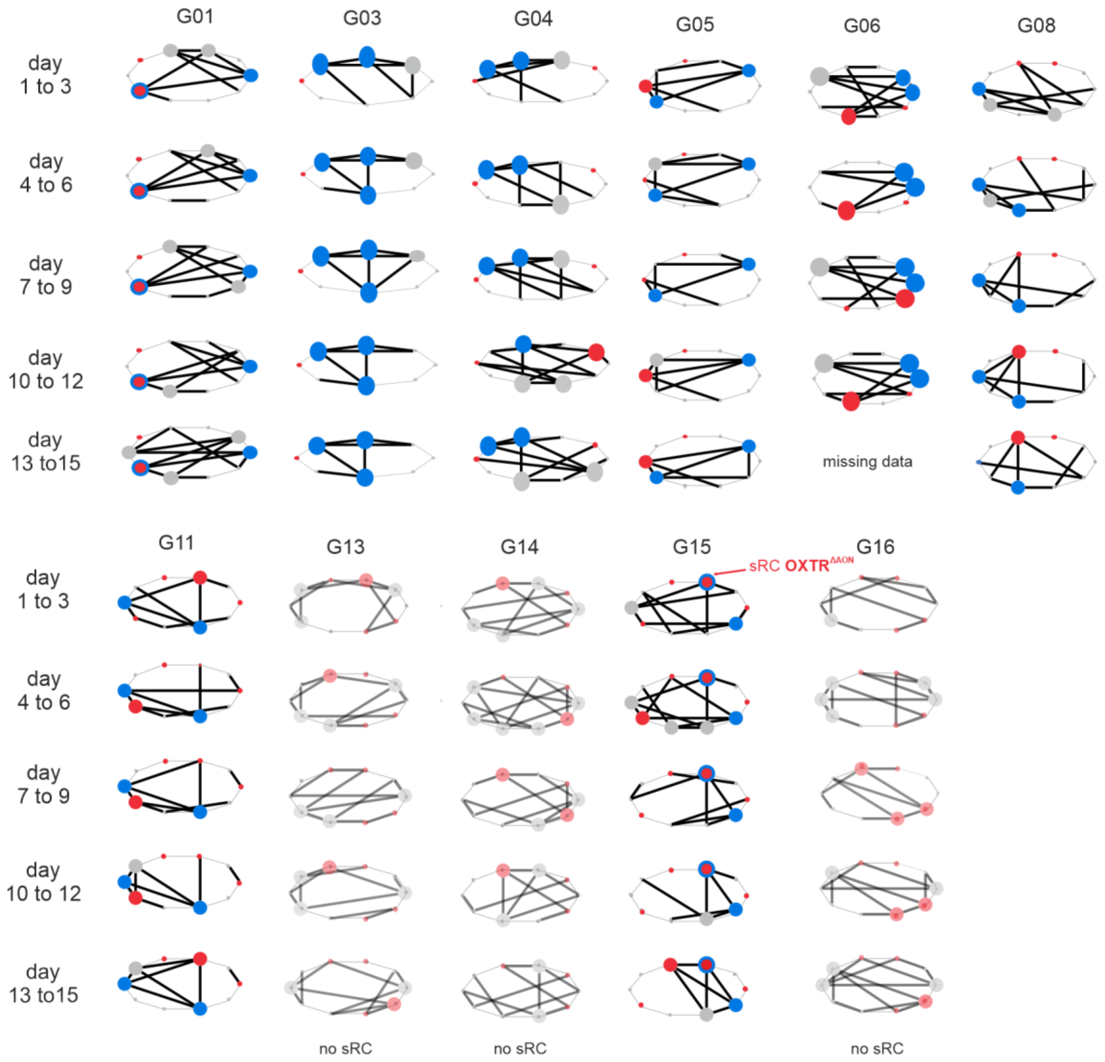


**Supplementary Fig. 5: Rich-club dynamics across different NoSeMaze groups.**
Rich-club (RC) dynamics are shown for 11 groups (G01, G03–06, G08, G11, G13–G16) across five consecutive 3-day intervals, displayed in horizontal columns. The remaining groups (G02, G07, G10, G12, G17) are displayed in Fig 4c of the main text. Each node represents an individual mouse. Edges reflect the total number of interactions between two mice, with the black edges representing the pruned networks using the mutual nearest-neighbor algorithm (see Methods). Stable RC (sRC) members, defined as nodes maintaining RC membership in ≥80% of time bins, are shown in bold blue; transient RC members in bold gray. OXTR$^{\Delta AON}$ mice are indicated in red. Groups are divided into older (top) and younger (bottom) sub-cohorts. Groups lacking stable RC structure (G13, G14, G16) are shown with light transparency. G06 has missing data for days 13–15. The consistent presence or absence of sRC members across time reflects inter-group variability in rich-club emergence and stability.

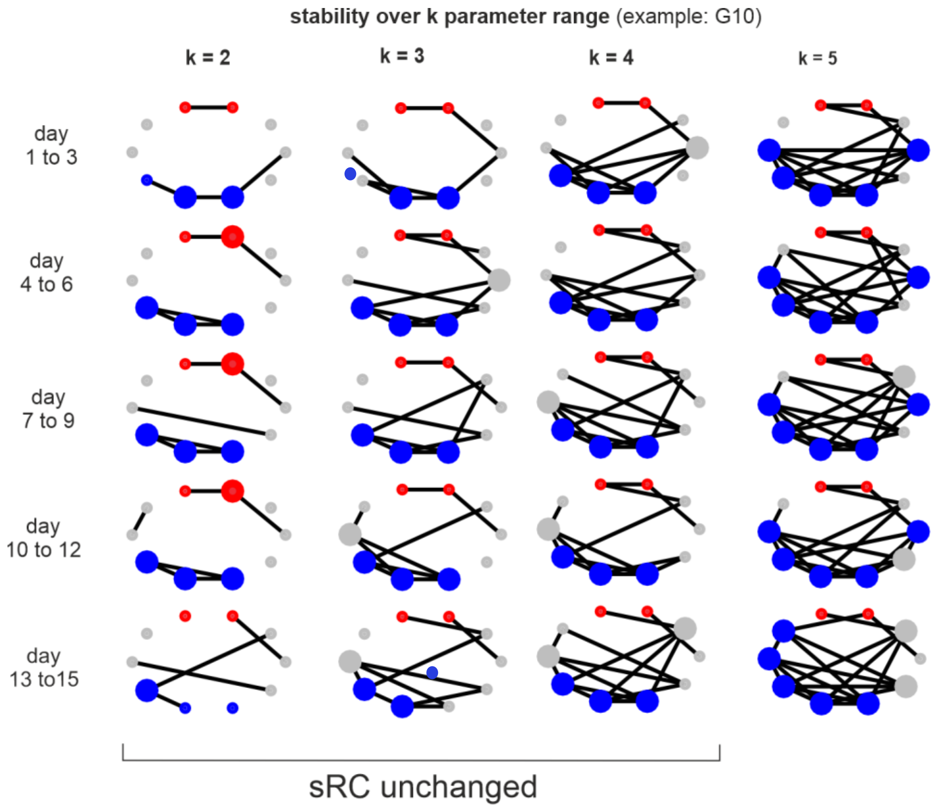


**Supplementary Fig. 6: Robustness of stable rich-club detection across different mutual nearest-neighbor pruning levels.** Rich-club (RC) structure for group G10 is shown across five 3-day windows and four mutual nearest-neighbor pruning levels (*k* = 2–5). Each node represents an individual mouse. Edges represent social ties that survive pruning, i.e., pairs of animals that are mutually ranked within each other's top *k* most frequent interaction partners (see Methods). RC members identified at each pruning level are shown as large nodes; non-members as small nodes. Stable RC members (present in ≥80% of time bins) are colored blue. $OXTR^{\Delta AON}$ mice are shown in red. As *k* increases, more edges are retained, but the core stable RC structure remains consistent, demonstrating robustness of RC detection to pruning stringency.

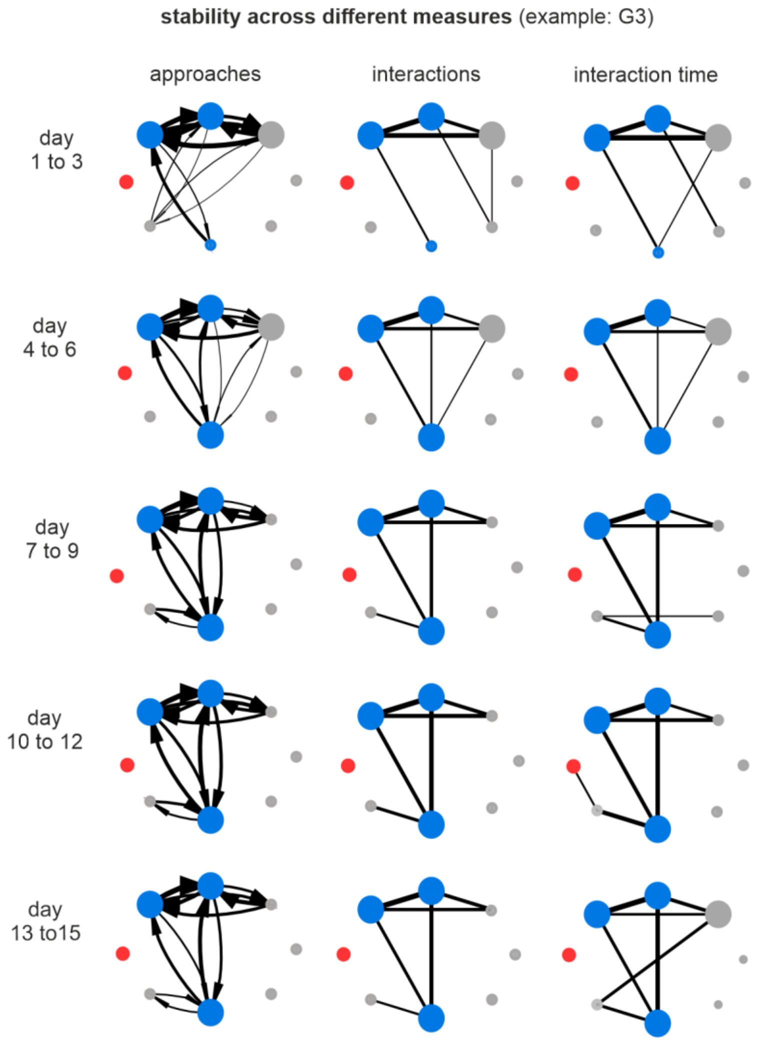


**Supplementary Fig. 7: Consistency of stable rich-club detection across different interaction metrics for social network construction.**

Rich-club (RC) structure for group G3 is shown across five 3-day windows and three network types based on different behavioral measures: approaches (left), number of interactions (middle), and interaction durations (right). Each node represents an individual mouse. Edges represent social ties that survive mutual nearest-neighbor pruning (k = 3), i.e., pairs of animals that are mutually ranked within each other's top three most frequent interaction partners (see Methods). RC members identified for each measure are shown as large nodes, non-members as small nodes. Stable RC members (present in ≥80% of time bins) are colored blue. $OXTR^{\Delta AON}$ mice are shown in red. Despite variation in the behavioral metric used for social network construction, the core RC structure remains consistent across network types, demonstrating the robustness of RC detection to different definitions of social interaction.

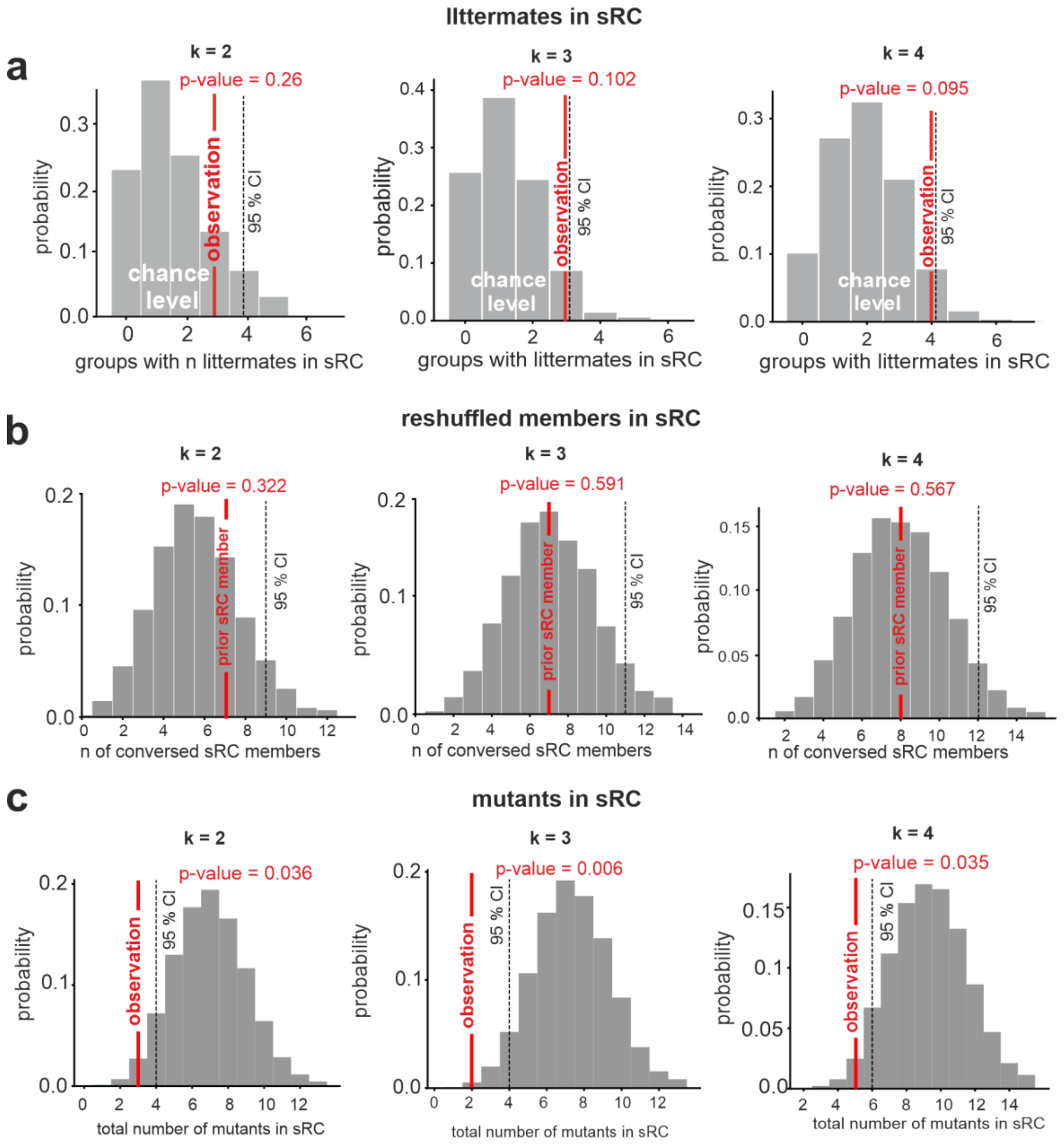


**Supplementary Fig. 8: Robustness of stable rich-club composition across different pruning parameters, assessed via permutation tests.**

**a)** Observed number of groups containing littermates within the same stable rich-club (sRC) compared to a null distribution generated by randomly shuffling group member identities (10,000 iterations). The observed values (red lines) fall within the 95% confidence intervals of the null model (gray distributions), indicating that shared family background does not significantly influence sRC formation ($p > 0.05$ for all *k* values. **b)** Number of conserved sRC members across reshuffled group contexts. The observed values (red lines) represent the number of prior sRC members who rejoined a sRC after random reassignment of group membership. Across all pruning levels (*k* = 2–4), the observed values fall within the null model distribution, suggesting that sRC membership is not an intrinsic or persistent individual trait, but rather emerges from the specific social dynamics of each group. **c)** Observed total number of $OXTR^{\Delta AON}$ mice within sRCs compared to a null distribution generated by random assignment. Across all pruning levels, observed frequencies (red lines) were significantly lower than expected by chance (all $p < 0.05$), indicating that $OXTR^{\Delta AON}$ mice are consistently underrepresented in stable rich-clubs.

For details on the three permutation procedures, see Methods.

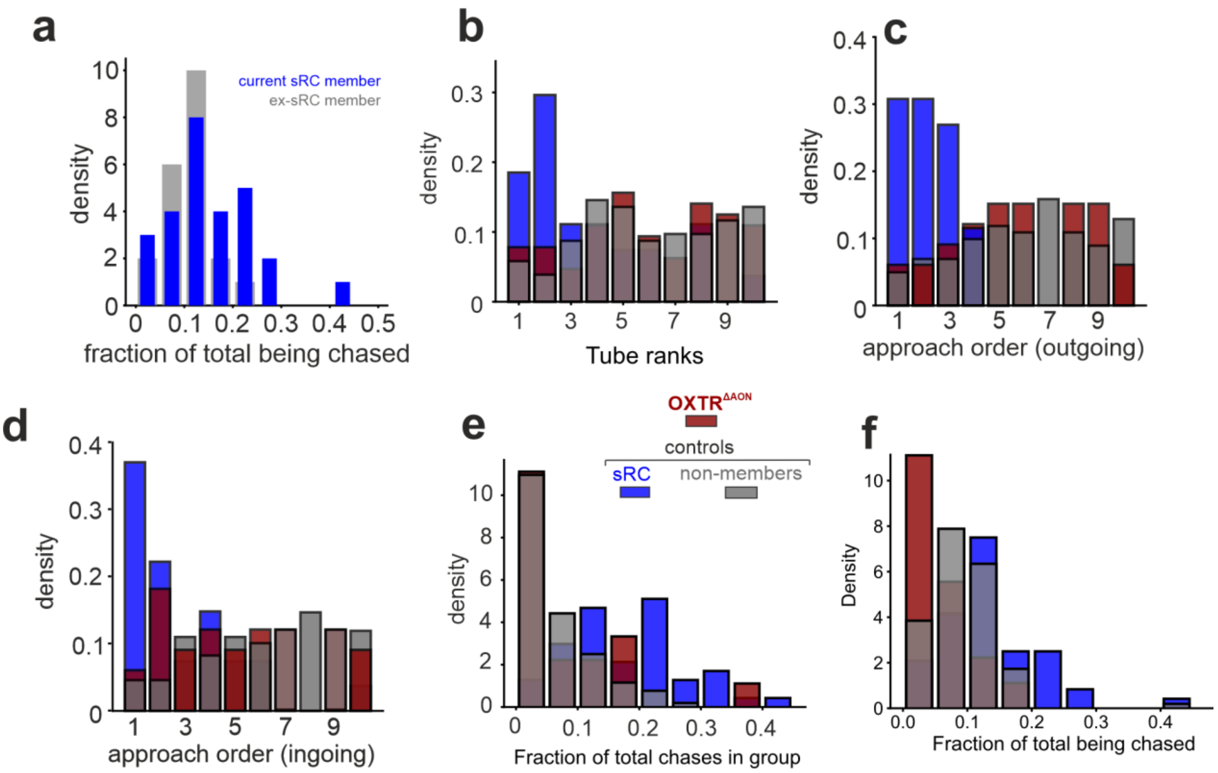


**Supplementary Fig. 9: Chasing and approach patterns across genotypes and stable rich-club (sRC) membership. a)** Fraction of total chases received by current sRC members (blue) and the same individuals when not in the sRC (ex-sRC members, gray). Being part of the sRC did not significantly change the distribution of incoming chases (p-value = 0.07). **b-f)** Distribution of behavioral measures of mice subdivided into three groups: sRC-members (blue), non-members (gray), and OXTR$^{\Delta AON}$ mice (red). sRC and non-members here only show control mice, so that no animal is represented more than once across groups. **b-d)** Normalized density distributions of within-group behavioral ranks (1 = highest activity) based on: **b)** tube-test dominance rank (competition outcomes), **c)** number of approaches initiated, and **d)** number of approaches received. **e)** Fraction of total chases initiated by mice across the three groups. **f)** Fraction of total chases received across the three groups. All densities are normalized (area under each curve = 1) to allow comparison across groups with unequal sample sizes. Significant differences between the distribution of sRC members (blue) and both non-members (gray) and OXTR$^{\Delta AON}$ mice (red) were observed in all cases ($p < 0.001$), while no differences were detected between non-members and OXTR$^{\Delta AON}$ mice.

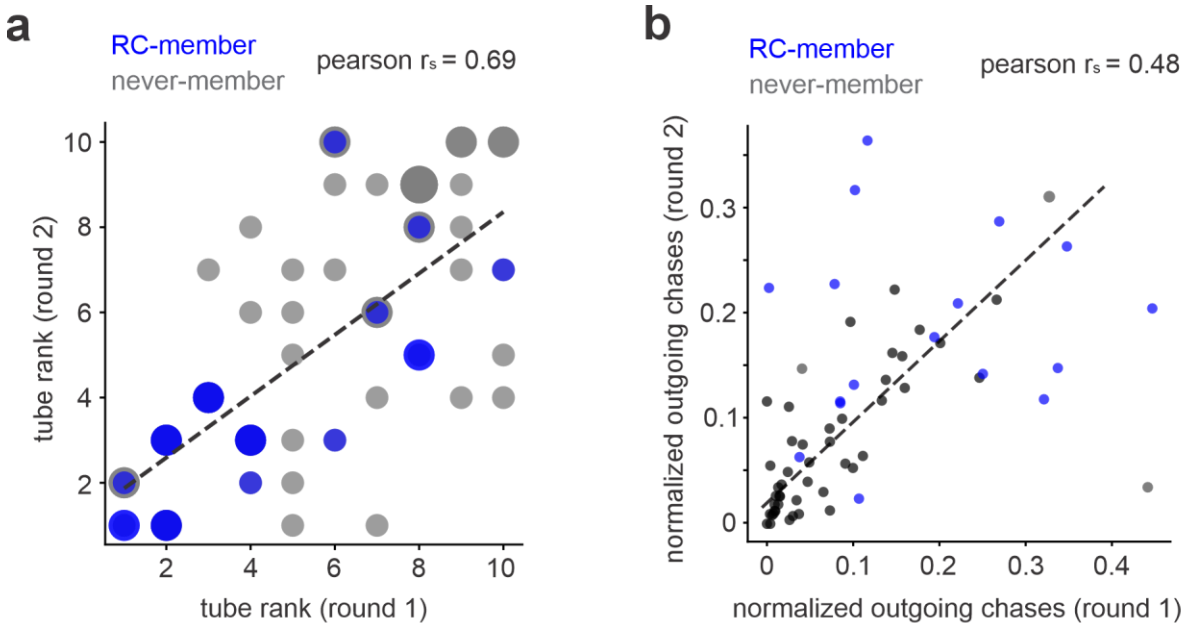


**Supplementary Fig. 10: Stability of tube-test rank and chasing behavior across rounds.**
**a)** Tube-test rank of mice in their first appearance in a NoSeMaze group plotted against their rank after reshuffling into a different group at a later round. Dot size reflects the frequency of a given value combination across rounds. While a clear overall correlation is observed, no significant difference was found between correlations of sRC members and non-members.
**b**) Fraction of initiated chases in the first group assignment compared to the second after reshuffling. A moderate overall correlation is observed, independent of sRC membership.

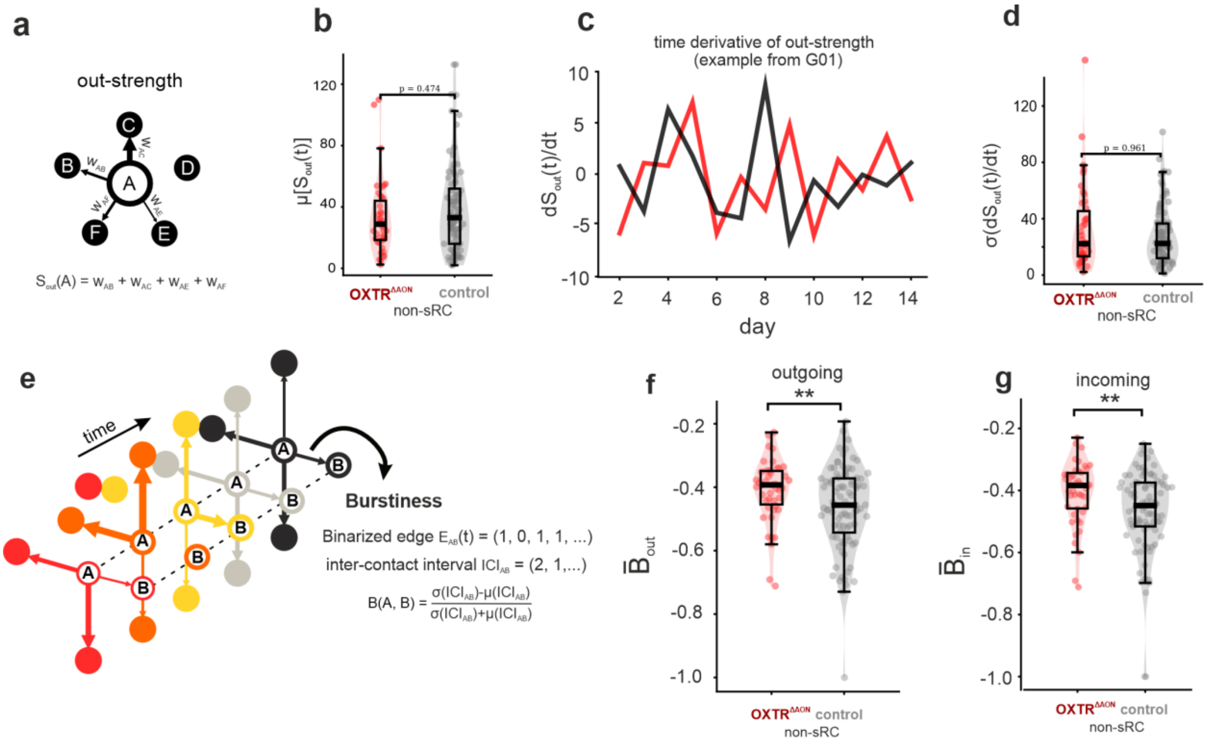


**Supplementary Figure 11: Outgoing strength and burstiness as measures of social interaction dynamics.**

**a)** Schematic illustrating outgoing strength ($S_{out}$), calculated as the sum of weighted edges from a node to all others. **b)** Mean $S_{out}$ did not differ significantly between OXTR$^{\Delta AON}$ mice and control non-members of the sRC. **c)** Example time derivatives of $S_{out}$ from an OXTR$^{\Delta AON}$ (red) and a control (dark gray) mouse from the same group. **d)** Standard deviation of the time derivative of $S_{out}$, used to assess fluctuations. Because $S_{out}$ reflects overall social output of a given mouse, it can be viewed as a proxy for motivation to initiate social interactions. The absence of differences between OXTR$^{\Delta AON}$ mice and control non-members indicates that OXTR$^{\Delta AON}$ mice did not show impaired motivation during the experiment. **e)** Schematic illustrating burstiness (B), computed using a modified coefficient of variation ($B = \frac{\sigma-\mu}{\sigma+\mu}$), where μ and σ are the mean and standard deviation of inter-contact intervals obtained from all possible edges of a given node (see Methods). Node-level burstiness was defined as the average burstiness across all edges connected to that node. As this metric requires a binary graph, we applied a graph cut at $k$ = 7 to avoid oversparsification (see main text). **f, g)** Outgoing and incoming averaged burstiness, respectively. OXTR$^{\Delta AON}$ mice displayed significantly more bursty interaction patterns both from and towards them compared to controls (permutation test on the median, 10,000 iterations). Overall, burstiness values for OXTR$^{\Delta AON}$ mice were closer to 0, indicating more Poisson-like connectivity patterns than in their neurotypical peers.

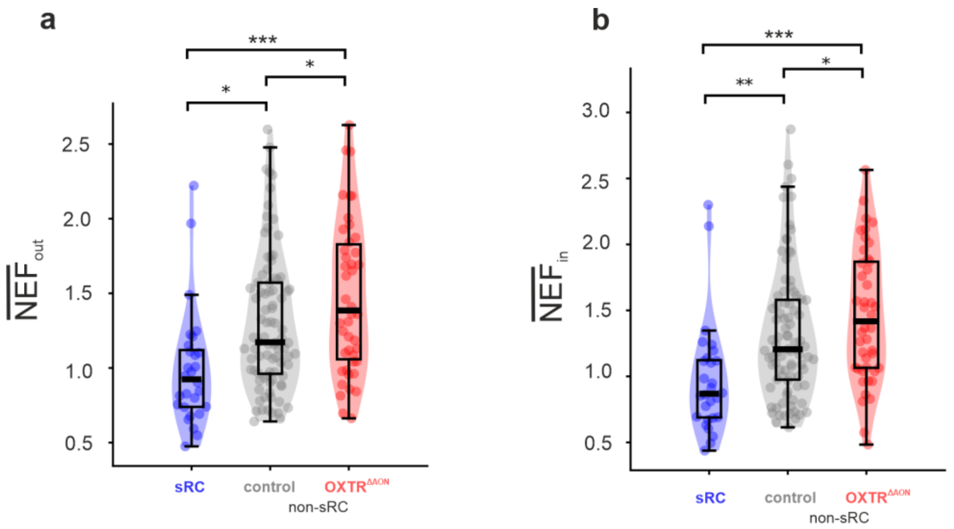


**Supplementary Figure 12: Stability of sRC interactions in time.**

Behavioral fluctuation metric (NEF) plotted for three groups: sRC members (blue), non-members (gray), and $OXTR^{\Delta AON}$ mice (red). sRC and non-members only show control mice, so that no animal is represented more than once across groups. Averaged NEF values are shown for the outgoing **a)** and ingoing **b)** direction, respectively. sRC members showed significantly lower NEF values than non-members or $OXTR^{\Delta AON}$ mice in both the outgoing (initiating) and incoming (receiving) direction (permutation test on the median, 10,000 iterations), indicating more consistent and regular social interactions. The strongest contract was observed between sRC members and $OXTR^{\Delta AON}$ mice ($p < 0.001$).

| Mouse RFID | group | repetition | age | genotype | weight | Mouse RFID | group | repetition | age | genotype | weight |
|---|---|---|---|---|---|---|---|---|---|---|---|
| 0007AC2B02 | 1 | 1 | 55 | WT | 33.59 | 0007CB0A48 | 12 | 1 | 18 | WT | 28.38 |
| 0007CB0942 | 1 | 1 | 55 | WT | 34.36 | 0007CB0AD4 | 12 | 1 | 16 | WT | 23.28 |
| 0007CB44CE | 1 | 1 | 55 | WT | 25.92 | 0007CB2ED2 | 12 | 1 | 16 | WT | 26.51 |
| 0007CB7051 | 1 | 1 | 56 | WT | 32.58 | 0007CB321B | 12 | 1 | 18 | WT | 26.61 |
| 0007CD6778 | 1 | 1 | 57 | $OXTR^{\Delta AON}$ | 36.42 | 0007CB39C9 | 12 | 1 | 18 | $OXTR^{\Delta AON}$ | 23.49 |
| 0007CD765E | 1 | 1 | 57 | WT | 31.68 | 0007CB464C | 12 | 1 | 16 | $OXTR^{\Delta AON}$ | 27.25 |
| 0007CD77C5 | 1 | 1 | 57 | $OXTR^{\Delta AON}$ | 32.71 | 0007F2B770 | 12 | 1 | 17 | $OXTR^{\Delta AON}$ | 27.57 |
| 0007CDADED | 1 | 1 | 55 | WT | 36.89 | 0007F2B957 | 12 | 1 | 18 | $OXTR^{\Delta AON}$ | 28.45 |
| 0007CDC443 | 1 | 1 | 55 | WT | 36.46 | 0007F2BA9B | 12 | 1 | 18 | WT | 26.04 |
| 0007CDDBAF | 1 | 1 | 57 | WT | 31.46 | 0007F2CA05 | 12 | 1 | 18 | $OXTR^{\Delta AON}$ | 25.31 |
| 0007CB2224 | 2 | 1 | 58 | WT | 35.99 | 0007CA3B1C | 13 | 1 | 17 | WT | 24.43 |
| 0007CB3491 | 2 | 1 | 56 | WT | 33.91 | 0007CB1E1F | 13 | 1 | 18 | WT | 28.15 |
| 0007CB4837 | 2 | 1 | 56 | $OXTR^{\Delta AON}$ | 40.03 | 0007CB2183 | 13 | 1 | 17 | $OXTR^{\Delta AON}$ | 25.32 |
| 0007CB6AB4 | 2 | 1 | 56 | WT | 27.19 | 0007CB2277 | 13 | 1 | 18 | $OXTR^{\Delta AON}$ | 26.24 |
| 0007CD5116 | 2 | 1 | 57 | WT | 36.69 | 0007CB3D9B | 13 | 1 | 18 | WT | 24.12 |
| 0007CDE223 | 2 | 1 | 75 | WT | 33.16 | 0007CB3F5E | 13 | 1 | 18 | WT | 25.26 |
| 0007CDE4F0 | 2 | 1 | 58 | $OXTR^{\Delta AON}$ | 30.71 | 0007CB6BC3 | 13 | 1 | 16 | WT | 23.81 |
| 0007CDEAA7 | 2 | 1 | 57 | WT | 36.04 | 0007F2945F | 13 | 1 | 18 | WT | 23.81 |
| 0007CDEC0B | 2 | 1 | 75 | WT | 32.32 | 0007F2B8A6 | 13 | 1 | 17 | $OXTR^{\Delta AON}$ | 24.56 |
| 0007CDEEE2 | 2 | 1 | 75 | WT | 32.92 | 0007F2E455 | 13 | 1 | 18 | $OXTR^{\Delta AON}$ | 24.72 |
| 0007CB0942 | 3 | 2 | 59 | WT | 34.47 | 0007CB0CDD | 14 | 1 | 16 | WT | 23.82 |
| 0007CD38E5 | 3 | 1 | 59 | WT | 35.51 | 0007CB0F67 | 14 | 1 | 16 | $OXTR^{\Delta AON}$ | 27.03 |
| 0007CD7FB2 | 3 | 1 | 63 | $OXTR^{\Delta AON}$ | 34.07 | 0007CB48B4 | 14 | 1 | 16 | WT | 27.09 |
| 0007CD81D3 | 3 | 1 | 59 | WT | 37.51 | 0007CB6DFA | 14 | 1 | 16 | $OXTR^{\Delta AON}$ | 24.81 |
| 0007CD86B3 | 3 | 1 | 78 | WT | 32.71 | 0007F29741 | 14 | 1 | 17 | WT | 25.78 |
| 0007CDAD6B | 3 | 1 | 78 | WT | 28.63 | 0007F2B88A | 14 | 1 | 17 | WT | 24.38 |
| 0007CDC395 | 3 | 1 | 63 | $OXTR^{\Delta AON}$ | 34.19 | 0007F2C687 | 14 | 1 | 18 | WT | 26.08 |
| 0007CDC91B | 3 | 1 | 78 | WT | 31.69 | 0007F2EEFA | 14 | 1 | 17 | WT | 24.78 |
| 0007CDDBAF | 3 | 2 | 61 | WT | 32.41 | 0007F2F26A | 14 | 1 | 18 | WT | 25.74 |
| 0007CE0327 | 3 | 1 | 78 | WT | 29.83 | 0007F2F283 | 14 | 1 | 16 | $OXTR^{\Delta AON}$ | 24.07 |
| 0007ABF555 | 4 | 1 | 62 | $OXTR^{\Delta AON}$ | 32.48 | 0007CA3A72 | 15 | 2 | 22 | $OXTR^{\Delta AON}$ | 26.26 |
| 0007AC2B02 | 4 | 2 | 60 | WT | 33.03 | 0007CB0B74 | 15 | 2 | 22 | WT | 27.02 |
| 0007ACA781 | 4 | 1 | 79 | WT | 32.69 | 0007CB0DBC | 15 | 2 | 22 | $OXTR^{\Delta AON}$ | 28.11 |
| 0007CB2224 | 4 | 2 | 62 | WT | 35.46 | 0007CB2407 | 15 | 2 | 22 | $OXTR^{\Delta AON}$ | 24.43 |
| 0007CB7051 | 4 | 2 | 61 | WT | 32.78 | 0007F2B3FF | 15 | 2 | 22 | WT | 26.18 |
| 0007CD6778 | 4 | 2 | 62 | $OXTR^{\Delta AON}$ | 38.56 | 0007F2B59E | 15 | 2 | 20 | WT | 28.52 |
| 0007CD765E | 4 | 2 | 62 | WT | 30.89 | 0007F2B8A2 | 15 | 2 | 22 | $OXTR^{\Delta AON}$ | 26.42 |
| 0007CDADED | 4 | 2 | 60 | WT | 37.41 | 0007F2B9FC | 15 | 2 | 22 | WT | 26.52 |
| 0007CDC443 | 4 | 2 | 60 | WT | 37.22 | 0007F2E37E | 15 | 2 | 22 | WT | 28.04 |
| 0007CDEAA7 | 4 | 2 | 60 | WT | 35.07 | 0007F2E3D4 | 15 | 2 | 22 | WT | 26.56 |
| 0007CB0942 | 5 | 3 | 62 | WT | 34.51 | 0007CA3B1C | 16 | 2 | 21 | WT | 26.49 |
| 0007CB3491 | 5 | 2 | 62 | WT | 32.79 | 0007CB1E1F | 16 | 2 | 22 | WT | 28.75 |
| 0007CB4837 | 5 | 2 | 62 | $OXTR^{\Delta AON}$ | 36.62 | 0007CB2183 | 16 | 2 | 21 | $OXTR^{\Delta AON}$ | 25.83 |
| 0007CD72E3 | 5 | 1 | 81 | WT | 35.94 | 0007CB2277 | 16 | 2 | 22 | $OXTR^{\Delta AON}$ | 27.41 |
| 0007CD77C5 | 5 | 2 | 64 | $OXTR^{\Delta AON}$ | 32.59 | 0007CB3D9B | 16 | 2 | 22 | WT | 25.72 |
| 0007CDDBAF | 5 | 3 | 64 | WT | 31.31 | 0007CB3F5E | 16 | 2 | 22 | WT | 26.67 |
| 0007CDE223 | 5 | 2 | 81 | WT | 31.39 | 0007CB6BC3 | 16 | 2 | 20 | WT | 26.87 |
| 0007CDEC0B | 5 | 2 | 81 | WT | 31.29 | 0007F2945F | 16 | 2 | 22 | WT | 24.34 |
| 0007CDEEE2 | 5 | 2 | 81 | WT | 31.82 | 0007F2B8A6 | 16 | 2 | 21 | $OXTR^{\Delta AON}$ | 26.76 |
| 0007CE0327 | 5 | 2 | 81 | WT | 28.41 | 0007F2E455 | 16 | 2 | 22 | $OXTR^{\Delta AON}$ | 25.24 |
| 0007CB6AB4 | 6 | 2 | 73 | WT | 29.18 | 0007CB0CDD | 17 | 2 | 20 | WT | 25.19 |
| 0007CB7051 | 6 | 3 | 74 | WT | 33.63 | 0007CB0F67 | 17 | 2 | 20 | $OXTR^{\Delta AON}$ | 29.21 |
| 0007CD5116 | 6 | 3 | 74 | WT | 31.61 | 0007CB48B4 | 17 | 2 | 20 | WT | 26.15 |
| 0007CD765E | 6 | 3 | 75 | WT | 31.39 | 0007CB6DFA | 17 | 2 | 20 | $OXTR^{\Delta AON}$ | 25.86 |
| 0007CD86B3 | 6 | 3 | 91 | WT | 32.58 | 0007F29741 | 17 | 2 | 21 | WT | 25.49 |
| 0007CDAD6B | 6 | 2 | 91 | WT | 27.82 | 0007F2B88A | 17 | 2 | 21 | WT | 25.22 |
| 0007CDADED | 6 | 3 | 73 | WT | 36.98 | 0007F2C687 | 17 | 2 | 22 | WT | 25.96 |
| 0007CDC395 | 6 | 2 | 77 | $OXTR^{\Delta AON}$ | 32.67 | 0007F2EEFA | 17 | 2 | 21 | WT | 25.61 |
| 0007CDC443 | 6 | 3 | 73 | WT | 34.07 | 0007F2F26A | 17 | 2 | 22 | WT | 26.72 |
| 0007CDE4F0 | 6 | 2 | 75 | $OXTR^{\Delta AON}$ | 30.01 | 0007F2F283 | 17 | 2 | 20 | $OXTR^{\Delta AON}$ | 25.74 |
| 0007ABF555 | 7 | 2 | 68 | $OXTR^{\Delta AON}$ | 31.61 | 0007CA3B1C | 18 | 3 | 26 | WT | 27.78 |
| 0007ACA781 | 7 | 2 | 85 | WT | 31.38 | 0007CB0AD4 | 18 | 3 | 25 | WT | 23.92 |
| 0007CB2224 | 7 | 3 | 68 | WT | 35.72 | 0007CB0F67 | 18 | 3 | 25 | $OXTR^{\Delta AON}$ | 29.38 |
| 0007CD38E5 | 7 | 2 | 66 | WT | 34.85 | 0007CB2183 | 18 | 3 | 26 | $OXTR^{\Delta AON}$ | 26.82 |
| 0007CD72E3 | 7 | 2 | 85 | WT | 34.39 | 0007CB2407 | 18 | 3 | 27 | $OXTR^{\Delta AON}$ | 25.84 |
| 0007CD7FB2 | 7 | 2 | 70 | $OXTR^{\Delta AON}$ | 33.79 | 0007CB3F5E | 18 | 3 | 27 | WT | 29.41 |
| 0007CD81D3 | 7 | 2 | 66 | WT | 34.29 | 0007CB48B4 | 18 | 3 | 25 | WT | 27.93 |
| 0007CDEAA7 | 7 | 3 | 66 | WT | 34.37 | 0007F2BA9B | 18 | 3 | 27 | WT | 28.36 |
| 0007CDEEE2 | 7 | 3 | 85 | WT | 30.45 | 0007F2E3D4 | 18 | 3 | 27 | WT | 28.97 |
| 0007CE0327 | 7 | 3 | 85 | WT | 29.61 | 0007F2E455 | 18 | 3 | 27 | $OXTR^{\Delta AON}$ | 26.29 |
| 0007CA3A3A | 8 | 1 | 68 | WT | 25.47 | 0007CB0AD4 | 19 | 2 | 20 | WT | 22.81 |
| 0007CB0942 | 8 | 4 | 68 | WT | 34.24 | 0007CB2ED2 | 19 | 2 | 20 | WT | 30.42 |
| 0007CB30A6 | 8 | 1 | 72 | $OXTR^{\Delta AON}$ | 30.41 | 0007CB321B | 19 | 2 | 22 | WT | 26.19 |
| 0007CB486D | 8 | 1 | 71 | $OXTR^{\Delta AON}$ | 31.92 | 0007CB39C9 | 19 | 2 | 22 | $OXTR^{\Delta AON}$ | 26.18 |
| 0007CD5116 | 8 | 2 | 69 | WT | 32.67 | 0007F2B770 | 19 | 2 | 21 | $OXTR^{\Delta AON}$ | 27.57 |
| 0007CD86B3 | 8 | 2 | 87 | WT | 32.68 | 0007F2B957 | 19 | 2 | 22 | $OXTR^{\Delta AON}$ | 30.19 |
| 0007CDC91B | 8 | 2 | 87 | WT | 32.73 | 0007F2BA9B | 19 | 2 | 22 | WT | 26.18 |
| 0007CDDBAF | 8 | 4 | 70 | WT | 32.51 | 0007F2CA05 | 19 | 2 | 22 | $OXTR^{\Delta AON}$ | 25.88 |
| 0007CDE223 | 8 | 3 | 87 | WT | 33.11 | 0007F2E42C | 19 | 1 | 22 | WT | 30.91 |
| 0007CDEC0B | 8 | 3 | 87 | WT | 31.56 | 0007CB0DBC | 20 | 3 | 27 | $OXTR^{\Delta AON}$ | 30.82 |
| 0007CB2224 | 9 | 4 | 73 | WT | 36.15 | 0007CB1E1F | 20 | 3 | 27 | WT | 30.65 |
| 0007CB2302 | 9 | 1 | 71 | $OXTR^{\Delta AON}$ | 28.67 | 0007CB321B | 20 | 3 | 27 | WT | 28.21 |

| Mouse RFID | group | repetition | age | genotype | weight | Mouse RFID | group | repetition | age | genotype | weight |
|---|---|---|---|---|---|---|---|---|---|---|---|
| 0007CB3713 | 9 | 1 | 71 | WT | 32.15 | 0007F29741 | 20 | 3 | 26 | WT | 27.97 |
| 0007CB40DE | 9 | 1 | 71 | WT | 25.89 | 0007F2B770 | 20 | 3 | 26 | $OXTR^{\Delta AON}$ | 29.14 |
| 0007CB47A6 | 9 | 1 | 72 | WT | 36.48 | 0007F2B9FC | 20 | 3 | 27 | WT | 27.96 |
| 0007CB486D | 9 | 2 | 74 | $OXTR^{\Delta AON}$ | 30.16 | 0007F2CA05 | 20 | 3 | 27 | $OXTR^{\Delta AON}$ | 26.27 |
| 0007CDEC0B | 9 | 4 | 90 | WT | 31.29 | 0007F2E37E | 20 | 3 | 27 | WT | 29.92 |
| 0007CDEEE2 | 9 | 4 | 90 | WT | 31.51 | 0007F2EEFA | 20 | 3 | 26 | WT | 28.03 |
| 0007CE0327 | 9 | 4 | 90 | WT | 29.14 | 0007F2F283 | 20 | 3 | 25 | $OXTR^{\Delta AON}$ | 27.64 |
| 0007CA3A3A | 10 | 2 | 75 | WT | 25.77 | 0007CA3A72 | 21 | 3 | 27 | $OXTR^{\Delta AON}$ | 27.51 |
| 0007CB0942 | 10 | 5 | 75 | WT | 35.77 | 0007CB2277 | 21 | 3 | 27 | $OXTR^{\Delta AON}$ | 29.32 |
| 0007CB2302 | 10 | 2 | 75 | $OXTR^{\Delta AON}$ | 28.45 | 0007CB464C | 21 | 3 | 25 | $OXTR^{\Delta AON}$ | 28.02 |
| 0007CB30A6 | 10 | 2 | 79 | $OXTR^{\Delta AON}$ | 29.28 | 0007F2B3FF | 21 | 3 | 27 | WT | 27.05 |
| 0007CB3713 | 10 | 2 | 75 | WT | 31.29 | 0007F2B59E | 21 | 3 | 25 | WT | 31.52 |
| 0007CB47A6 | 10 | 2 | 76 | WT | 35.97 | 0007F2B88A | 21 | 3 | 26 | WT | 28.08 |
| 0007CDDBAF | 10 | 5 | 77 | WT | 31.39 | 0007F2B8A6 | 21 | 3 | 26 | $OXTR^{\Delta AON}$ | 28.69 |
| 0007CDE223 | 10 | 4 | 94 | WT | 32.49 | 0007F2B957 | 21 | 3 | 27 | $OXTR^{\Delta AON}$ | 31.32 |
| 0007CDEC0B | 10 | 5 | 94 | WT | 30.08 | 0007F2C687 | 21 | 3 | 27 | WT | 27.81 |
| 0007CE0327 | 10 | 5 | 94 | WT | 28.62 | 0007F2F26A | 21 | 3 | 27 | WT | 28.52 |
| 0007CA3A72 | 11 | 1 | 18 | $OXTR^{\Delta AON}$ | 26.68 | | | | | | |
| 0007CB0B74 | 11 | 1 | 18 | WT | 27.92 | | | | | | |
| 0007CB0DBC | 11 | 1 | 18 | $OXTR^{\Delta AON}$ | 28.07 | | | | | | |
| 0007CB2407 | 11 | 1 | 18 | $OXTR^{\Delta AON}$ | 24.12 | | | | | | |
| 0007F2B3FF | 11 | 1 | 18 | WT | 25.38 | | | | | | |
| 0007F2B59E | 11 | 1 | 16 | WT | 28.07 | | | | | | |
| 0007F2B8A2 | 11 | 1 | 18 | $OXTR^{\Delta AON}$ | 25.69 | | | | | | |
| 0007F2B9FC | 11 | 1 | 18 | WT | 25.71 | | | | | | |
| 0007F2E37E | 11 | 1 | 18 | WT | 26.1 | | | | | | |
| 0007F2E3D4 | 11 | 1 | 18 | WT | 26.16 | | | | | | |

**Table S1: Overview of group compositions and animal information.**

*$OXTR^{\Delta AON}$, bilateral oxytocin receptor deletion in the AON pars centralis; RFID, radio-frequency identification; WT, wild type*

| group ID | n of mice | age (median, SD, in weeks) | tube data | stimulus-outcome learning data | video data |
|---|---|---|---|---|---|
| 1 | 10 | 55.5, 0.99 | acquired | acquired | acquired |
| 2 | 10 | 57.5, 8.79 | acquired | acquired | acquired |
| 3 | 10 | 63, 9.07 | acquired | acquired | acquired |
| 4 | 10 | 61.5, 5.77 | acquired | acquired | acquired |
| 5 | 10 | 72.5, 9.62 | acquired | acquired | acquired |
| 6 | 10 | 74.5, 7.17 | acquired | acquired | acquired |
| 7 | 10 | 69, 9.20 | acquired | acquired | acquired |
| 8 | 10 | 71.5, 9.03 | acquired | acquired | acquired |
| 9 | 9 | 73, 9.06 | acquired | acquired | insufficient quality |
| 10 | 10 | 76.5, 8.78 | acquired | acquired | acquired |
| 11 | 10 | 18, 0.63 | acquired | acquired | acquired |
| 12 | 10 | 18, 0.95 | acquired | missing | acquired |
| 13 | 10 | 18, 0.81 | acquired | missing | acquired |
| 14 | 10 | 16.5, 0.82 | acquired | missing | acquired |
| 15 | 10 | 22, 0.63 | acquired | missing | acquired |
| 16 | 10 | 22, 0.71 | missing | acquired | acquired |
| 17 | 10 | 20.5, 0.82 | acquired | acquired | acquired |
| 18 | 10 | 26.5, 0.92 | acquired | acquired | missing |
| 19 | 10 | 22, 0.88 | acquired | acquired | missing |
| 20 | 10 | 27, 0.84 | missing | acquired | missing |
| 21 | 10 | 27, 0.84 | missing | acquired | missing |

**Table S2: Overview of group characteristics and acquisition of behavioral data in the NoSeMaze.**
*ID, identity; NoSeMaze, non-invasive sensor rich maze; STD, standard deviation*

### Extended Methods

### Animal strains, husbandry and preparation

**Dyadic social interaction experiment.** To investigate the effects of oxytocin system modulation on dyadic social interaction behavior, we used two distinct male experimental cohorts targeting different aspects of the oxytocin pathway (cf. Fig. 1c-d). The first cohort involved a conditional, adult-specific deletion of the oxytocin receptor (OXTR) in the AON ($OXTR^{\Delta AON}$). This was achieved by bilateral injection of an AAV-Cre vector (*$rAAV_{1/2}$-CBA-Cre*) into the AON of $OXTR_{fl/fl}$ mice (B6.129(SJL)-$Oxtr^{tm1.1Wsy}$/J, RRID: IMSR_JAX:008471, Jackson Laboratory**;** n=6 mice) (see also *'Virus preparation and stereotactic surgery'*). These mice were compared to matched $OXTR_{fl/fl}$ mice injected with a control virus (*$rAAV_{1/2}$-CBA-dTomato*, n=6 mice). The second cohort involved optogenetic stimulation of oxytocin-releasing neurons in the paraventricular nucleus (PVN) of the hypothalamus. In this group (n = 6), Cre-dependent expression of channelrhodopsin-2 (ChR2) was achieved via AAV injection (*$rAAV_5$-DIO-hChR2(H134R)-mCherry*, Addgene #20297-AAV5) in male OXT-Cre mice (B6;129S-*$Oxt^{tm1.1(cre)Dolsn}$*/J, RRID:IMSR_JAX:024234, Jackson Laboratory), allowing temporally controlled enhancement of endogenous oxytocin release (see also *'Virus preparation and stereotactic surgery'*). Corresponding control sessions were conducted without optical stimulation, allowing each animal to serve as its own control. All mice were at least 16 weeks old and were maintained on a C57BL/6J background, backcrossed for more than 10 generations (Charles River, Sulzfeld)**.** Animals were single-housed under controlled environmental conditions (12:12 h light/dark cycle, 24 °C, 55% humidity), with ad libitum access to food and water.

**Virus preparation and stereotactic surgery.** Distinct viral constructs were used to manipulate oxytocin signaling in the two experimental cohorts**.** The procedure has been previously described in detail **(**Wolf et al., 2024**).** To induce conditional *OXTR* deletion in the first cohort, six mice received bilateral stereotactic injections of an adeno-associated virus carrying Cre recombinase (*$rAAV_{1/2}$-CBA-Cre*) into the AON pars centralis, while the remaining six animals received a control virus encoding only dTomato (*$rAAV_{1/2}$-CBA-dTomato*). In the second cohort, a Cre-dependent adeno-associated virus expressing channelrhodopsin-2 (*$rAAV_5$-DIO-hChR2(H134R)-mCherry*) was injected **to allow** optogenetic activation of OXT neurons in the paraventricular nucleus (PVN). All stereotactic surgeries were performed under isoflurane anesthesia (induction at 3–4%, maintenance at 1–2%) with pre- and post-operative analgesia (meloxicam, Metacam, Boehringer Ingelheim). Mice were at least 10 weeks old and maintained on a heating pad for the duration of the procedure. After securing the head in a stereotaxic frame (Kopf Instruments), the skull was leveled and a local anesthetic (lidocaine) was applied to the incision site. Virus was delivered using a pulled-glass micropipette attached to a nanoinjector (MO-10, Narishige). A total volume of 0.5 µl was infused at two sites per hemisphere. Coordinates for PVN injections were: 0.1 or 0.3 mm posterior to bregma, 1.0 mm lateral, 4.8 mm ventral from skull surface, delivered at a 10° angle relative to vertical. The optic fiber (FT-200-EMT with ceramic ferrule CFLC230-10, Thorlabs) for optogenetic activation of the PVN-OXT neurons was implanted according to the following coordinates relative to bregma (in mm): 0.2 posterior, 0.8 lateral, 4.3 ventral, at an angle of 10° to the vertical axis. For OXTR deletion in AON, injection coordinates were: 3.0 mm anterior to bregma, 0.7 and 1.2 mm lateral, and 3.5 mm ventral.

**NoSeMaze experiment.** 79 adult male homozygous is

**Ethics statement.** All procedures were in accordance with the National Institutes of Health Guide for the Care and Use of Laboratory Animals and the EU 2010/63 directive, and approved by the local animal welfare authority (Referat 35, Regierungspräsidium Karlsruhe, Karlsruhe, Germany).

**Sex.** This study focused on male mice, for whom social network dynamics are most robustly characterized [1], and for whom the NoSeMaze system was originally optimized. Including both sexes in the same experimental environment leads to mating behavior, which fundamentally alters social dynamics and network structure. Ongoing work is aimed at adapting the NoSeMaze for use with female cohorts, with the goal of extending these findings to female-only or mixed-sex social systems in future research.

**Immunohistochemistry.** Following completion of the experiments, animals were anesthetized and transcardially perfused for histological verification of viral expression. Brains were fixed in 4% paraformaldehyde for two weeks, and then coronally sectioned (50 µm) and stained with antibodies against OXT (anti-OXT (1:1000, mouse; kindly provided by Harold Gainer)), Cre (anti-Cre (1:1000, rabbit; Novagen, cat. n. 69050-3)), or GFP (anti-GFP (1:1000, chicken, Abcam, cat. n. 13970)) diluted in PBS containing 0.2% Triton X-100 (Merck) overnight at 4°C. The signals were visualized with ALEXA 488 secondary antibody (goat IgG, 1:1000, ThermoFisher Invitrogen, anti-mouse cat. n. A-11001, anti-rabbit cat. n. A-11008, or anti-chicken A-11039) respectively incubated for 2 hours at room temperature. Fluorescent signals were visualized using either a confocal microscope or a slide-scanning system, in accordance with standard immunohistochemistry protocols (see Wolf et al., 2024). Images were acquired with an Olympus Axio Imager 2 microscope and expression patterns were mapped onto a standard brain atlas.

### Free social interaction paradigm

Each mouse in the $OXTR^{\Delta AON}$ group and its corresponding control group participated in two interaction sessions. In contrast, mice in the optogenetic stimulation cohort completed four sessions: two with optogenetically induced oxytocin release and two interleaved control sessions. During control sessions, optical stimulation was prevented by blocking light transmission through insertion of a small non-transparent plastic disc at the junction (mating sleeve) between the ferrule from the patch cord and the optical fiber implant. To minimize fatigue and allow behavioral recovery, animals were given one week between each interaction sessions.

During each session, the experimental mouse was placed in a clean standard cage (under 70 lux ambient lighting) and introduced to an unfamiliar, adolescent, same-sex C57BL/6 (postnatal day 35-50). This conspecific was selected to minimize the likelihood of aggression or sexual behavior. The dyad was allowed to interact freely for 5 minutes. All sessions were recorded at 30 frames per second using a Sony FDR-X1000V camera. In sessions involving optogenetic stimulation, light pulses (60 pulses per train, 5 ms duration, 30 Hz) were delivered every 30 seconds throughout the interaction period to induce oxytocin release.

**Analysis of interaction videos.** All video recordings were trimmed to 8,092 frames (~4 minutes and 30 seconds) to standardize video length. Frame alignment and resolution were unified to avoid distortions during analysis. Behavioral annotation was conducted manually using BORIS (Behavioral Observation Research Interactive Software [2]), an open-source tool for labeling video-recorded behavior. A custom ethogram was used to classify behaviors into discrete states: anogenital sampling, nose-to-nose sampling, flank sampling, approach, pursuit, attack, climbing, digging, and self-grooming (cf. Figure 1a-b). All video labeling was blinded to the condition of the mouse (Cre- or dTomato- expression, or optogenetic or sham stimulation). Behaviors were manually labeled using keyboard shortcuts, and outputs were exported in CSV format for further analysis. Inactivity or behaviors unrelated to social interaction like climbing, digging, and self-grooming were post-hoc classified as 'idle' states. Pursuit was grouped with approach. No attacks were observed in any session. Subsequent analyses focused on five behavioral states: anogenital sampling, nose-to-nose sampling, flank sampling, approach, and idle. The three sampling behaviors were further grouped as 'interaction states' for the state-based analysis, but were kept separate in the transition analysis. Mean duration and frequency of interaction states, approach, and idle were compared using linear mixed-effect models (LME), with animal ID as a random effect to account for repeated measures. In the optogenetic release cohort, sham sessions were compared to oxytocin-release sessions; in the $OXTR^{\Delta AON}$ group, $OXTR^{\Delta AON}$ mice and control animals were compared.

To assess sequential behavioral patterns, transition matrices were computed for each animal based on the annotated behavioral state sequences. A transition was defined as a change from one behavioral state (e.g., approach) to another (e.g., nose-to-nose sampling) across consecutive time points. Each matrix represented the relative probability of transitioning from a given source state to a target state, normalized over all outgoing transitions. Matrices were calculated at the individual level and then averaged across animals within each experimental condition (control, $OXTR^{\Delta AON}$, sham, oxytocin release). Then, for each transition between behavioral states, LMEs were fit to compare either control vs. $OXTR^{\Delta AON}$ or sham vs. oxytocin release, using group (or session type) as a fixed effect and animal ID as a random effect with random slopes. Together with the average matrices for the four experimental conditions, the resulting *t*-statistics from the LMEs were visualized in matrix form to identify significant differences in transition probabilities between conditions. False discovery rate (FDR) correction was applied to account for multiple comparisons.

### NoSeMaze

**NoSeMaze setup.** The NoSeMaze is a complex, semi-naturalistic environment designed for continuous, long-term assessment of social behavior, social status, and reinforcement learning in freely interacting mouse groups without human interference [3]. The reinforcement learning module is constructed based on an open-source Python framework [4], The full documentation is available at https://github.com/KelschLAB/NoSeMaze. Four identical NoSeMaze units operated in parallel, each housing an independent social group of 9–10 mice. Each NoSeMaze unit consists of two connected zones: a housing arena, containing nesting material and free access to food, and an open-field arena where social interactions take place (cf. Fig. 2a). The arenas are linked by two tubes equipped with RFID readers at each end that allow for automated tracking of individual movements and interactions. Water is delivered exclusively via a stimulus-outcome learning module connected to the open-field arena, where animals initiate trials ad libitum to meet their water needs. This design ensures that animals regularly move between the arenas to access both food and water, thereby promoting naturalistic patterns of exploration and interaction. The open-field arena is continuously recorded using overhead infrared cameras, enabling 24/7 video monitoring of spontaneous group behavior throughout each experimental round. All animals learned to use the water lickport, remained healthy, and none were excluded due to injury or social incompatibility over the course of the experiment. Thus, the NoSeMaze provides a relatively peaceful environment with no overt aggression, supporting stable group coexistence.

**Experimental design and animal grouping.** A total of 79 male mice were used in the NoSeMaze experiments, divided into two age-defined subcohorts (cf. Fig. 2b; see also 'Animal Strains and

Preparation' and Supplementary Table S1). The older subcohort (n = 38) entered the NoSeMaze for the first time at 55 weeks of age, while the younger subcohort (n = 41) began participation at the age of 16 weeks. OXTR depletion was performed at least six weeks before the start of the experiments. Over the course of the study, animals were tested across a total of 21 NoSeMaze rounds, each lasting three weeks (see Supplementary Table S2). The majority of mice (n=75) participated in two or more rounds. During each NoSeMaze round, mice were housed in dynamically reconfigured social groups of 9–10 individuals for three weeks, with no human interference during the period of the three weeks round. To examine behavioral stability across changing social contexts, group compositions were reshuffled from one round to the next (cf. Fig. 2c). Mice returned to home cages in groups of four between NoSeMaze rounds, with regular regrouping to promote familiarity among potential future group members. The average interval between the start of two successive NoSeMaze rounds was 6.8 weeks (SD = 2.8 weeks). Prior to entering the NoSeMaze, all mice underwent two habituation sessions in the NoSeMaze, conducted in subgroups of 6–7 animals. Each habituation session lasted 16–20 hours and included one full dark cycle, allowing mice to acclimate to the environment and learn the basic functionality of the water lickport (for details on gradual acclimatization, see [3]*)*. To enable individual identification in the continuous video recordings, mice were marked using non-toxic fur bleaching before each round under light anesthesia (1 % isoflurane) (cf. Fig. 3c). These fur patterns were used to train identification networks in DeepLabCut for automated tracking.

**Quantification of individual reward-seeking and social behavioral metrics.** This study primarily focused on video-based analysis of spontaneous social behavior recorded continuously in the open-field arena of the NoSeMaze (see *'Social network analysis'* for details). In addition, behavioral tracking included automated assessment of reinforcement learning based on task performance and social dominance using RFID-based monitoring in the tubes (for detailed methodological description, see [3]).

In brief, mice obtained water through the go-no-go task at the stimulus-outcome learning module with the water lick port, which is connected to the open-field arena. Animals learned to discriminate between two odors (Decanal, Sigma Aldrich W236217, and Octanal, Sigma Aldrich W279706), of which one was rewarded (CS+) and one was not rewarded (CS−). Mice received a water drop in 100% of go trials (CS+) if they licked twice during odor presentation (from 0.5 to 2.5 s). In no-go trials (CS−), no reward was given, and licking more than once during odor presentation resulted in a six-seconds timeout. Odor-reward contingencies switched every three days. Task engagement, licking behavior, and individual water intake were tracked continuously and linked to each animal's RFID identity. Multiple reward-seeking metrics were extracted, including correct hit and rejection rates, pre-CS licking rates, and switch latencies after reversals (for details on calculation and stability of metrics, number of trials, and daytime of activity, see [3]*)*. All trials were used to calculate pre-CS licking, correct hit, and correct rejection rates. CS+ switch latency measured how quickly an animal increased licking in response to the newly rewarded stimulus (CS+) after a reversal. It was defined as the number of CS+ trials needed post-reversal for licking to reach at least 70% of the animal's pre-reversal CS+ rate. CS- switch latency captured how quickly the animal suppressed licking to the newly unrewarded stimulus (CS-), defined as the number of CS- trials needed for licking to fall below 50% of the pre-reversal CS+ rate. Pre-reversal rates were based on the last 150 CS+ trials, measured during the odor window (0.5–2.5 s). If no switch occurred, latency was set to the total number of available post-reversal trials. Both metrics reflect individualized behavioral adaptation, relative to each animal's own baseline.

Social rank was inferred from interactions occurring in the two tubes connecting the housing and open-field arenas. These tubes were equipped with RFID readers and represented natural bottlenecks that mice had to pass through to access food or water, creating opportunities for competitive and dominance-related interactions. Tube competitions were identified when two animals entered the same tube from opposite ends, resulting in one mouse pushing the other out. The dominant mouse was detected at both ends of the tube, while the subordinate mouse was detected twice at the same entry point, indicating retreat (cf. Fig. 5d). Chasing events were defined as two mice moving in the same direction through a tube in rapid succession, based on specific timing thresholds (i.e., <1.5 s at entry detector, <2s total transit time, cf. Fig. 5f). Dominance hierarchies were quantified for tube competitions using David's scores [5], a standard network-based metric that accounts for both direct wins and losses as well as the hierarchical position of the opponents [3]. In addition, individual social behavior was characterized by the proportion of initiated and received chases [3].

### Video recording of social interactions

**Video recording and processing.** During the NoSeMaze experiment, continuous 24/7 video recordings were conducted in the open-field arena to monitor spontaneous social interactions. A USB 2.0 camera (Aptina MT9V022, FLIR) equipped with a wide-angle lens (focal width: 1.67 mm) captured the arena from above (cf. Fig. 3a). Red LED light strips illuminated the social observation arena, ensuring adequate video quality during both light and dark phases. Videos were available for 17 groups (see Supplementary Table S2), resulting in a total of approximately 6,100 hours of continuous video data that were analyzed using DeepLabCut (DLC) [6]. Separate DLC models based on the ResNet-50 architecture were trained for each group to detect the different fur bleaching patterns in the video frames, enabling individual identification and analysis of interaction behavior (cf. Fig. 3c-d). Training datasets were compiled from video frames taken on days 1, 8, and 15 of the experiment. Frame selection was performed using DLC's integrated k-means algorithm, which automatically extracts the most dissimilar frames from periods when at least one mouse was clearly visible. Fur patterns were then manually labeled by trained experimenters, ensuring balanced representation of individuals in the training set. Experimenters were fully blind to the whether an animal was neurotypical or $OXTR^{\Delta AON}$. Models were refined iteratively by adding additional training frames until satisfactory prediction accuracy was reached on previously unseen videos (defined as <5-pixel test error with a likelihood p-cutoff of 0.8, and visually validated performance). After training, the entire video dataset was processed using the group-specific DLC models. The output – x-y coordinates and likelihood values for each detected mouse – was post-processed to improve accuracy. Predictions with a likelihood below 0.8 were discarded to reduce false positives. Additionally, a temporal filter was applied: mice had to be consistently detected in at least 66% of frames over continuous intervals of at least one second. For two animals in group 3, the models failed to reach sufficient prediction accuracy; these individuals were excluded from further video-based analyses. For group 9, the models failed to reach sufficient prediction accuracy in 4 out of 9 mice, so the entire group was excluded. Moreover, as tracking accuracy declined in the third week of each round due to fading of the fur patterns, data from the third week were excluded from all analyses.

**Video analysis.** First, the time each animal spent in the arena was quantified individually. Next, social interaction events were extracted based on spatial proximity: an interaction was defined as any instance where the distance between two mice was less than 10 cm for at least 1 second. Successive events separated by fewer than 5 frames were merged into a single interaction event. For each social interaction, we additionally assessed whether one mouse actively approached the other. To determine this, the smoothed trajectories of both animals were analyzed during the 4 seconds preceding the interaction. If the distance traveled by one mouse exceeded that of the other by a factor of 1.5 or more, it was classified as the approaching individual. If neither met this threshold – such as when both mice were moving in parallel or converging passively – no approach was assigned.

### Social network analysis

**Network construction and basic metrics.** By tracking all events occurring between each pair of animals on a daily basis, we obtained social networks of dyadic (pairwise) interactions. These networks were represented as square matrices, with off-diagonal entries indicating the number of events for each animal pair. Social interaction networks were undirected, as no distinction was made between interaction partners; accordingly, the matrices were symmetric, with the number of interactions from A to B equal to those from B to A. In contrast, approach networks were directed, capturing how animals initiated or received approaches, resulting in asymmetric matrices. Since each social interaction event had a duration (measured by the number of frames the animals remained within a 10 cm radius), we also constructed undirected networks of mean interaction time, representing the average duration each pair of mice interacted per day.

We first analyzed networks of interactions, approaches and interaction time aggregated over the first two weeks of the experiment. To assess group differences between OXTR$^{\Delta AON}$ and control animals, we calculated: (1) the total number of approaches initiated (outgoing) or received (ingoing) by each animal, (2) the total number of social interactions involving each animal, and (3) the average total time each animal spent interacting. These values were computed by summing or averaging the corresponding rows or columns of the aggregated matrices, depending on the respective metric and the symmetry of the matrix. Statistical differences between OXTR$^{\Delta AON}$ and control animals were assessed with permutation tests on the median (10,000 permutations).

**Graph pruning.** Beyond analyzing aggregated measures such as the total number of interactions or outgoing approaches, we aimed to investigate the presence of specific substructures within the social networks and how individual animals contributed to them. To this end, we represented the networks as graphs, where nodes corresponded to individual animals and edges reflected the number of events involving each animal pair. However, due to the high volume of daily interactions, these graphs were densely connected, potentially obscuring meaningful structure. Graph pruning was therefore necessary to reveal meaningful patterns. Since activity levels varied substantially across groups, standard pruning methods using fixed absolute or relative edge thresholds were impractical. Instead, we applied a mutual nearest-neighbor approach. Specifically, we retained an edge between two animals (A and B) only if they were among each other's top *k* interaction partners. That is, B had to be among A's *k* most frequent interaction partners, and A among B's. We tested several *k* values (see Supplementary Fig. 6 and 8), and all results shown in this article were obtained using *k* = 3. The procedure is visualized in Fig. 4a.

**Rich-clubs.** Having obtained sparser and more interpretable graphs, we next examined whether certain animals formed tightly interconnected subgroups. Specifically, we investigated the presence of *rich-clubs, defined as groups* of highly connected individuals (i.e., nodes with high degree) that are also densely connected to one another. Formally, a rich-club of degree *k* refers to the subset of nodes *v* with degree $deg(v) \geqslant k$ that are more interconnected than expected by chance. To assess this in weighted networks, we computed the weighted rich-club coefficient following the framework proposed by Alstott et al. (2014). In this approach, the observed weighted rich-club coefficient $\phi(k)$ is compared to that obtained from randomized control networks, $\phi_{rand}(k)$. Thus, the *normalized* weighted rich-club coefficient is defined as:

$$\phi_{norm}(k) = \frac{\phi(k)}{\phi_{rand}(k)} = \frac{\frac{C(k)}{F(k)}}{\frac{C_{rand}(k)}{F_{rand}(k)}}$$

where:

- $C(k)$ is the weighted connectedness of the club, defined as the sum of the weights of the edges between the rich-club nodes in the empirical network [7].
- $F(k)$ is the maximal possible weighted connectedness of the rich-club, defined as the sum of the weights of all edges connected to the rich nodes under the given assumptions.
- $C_{rand}(k)$ and $F_{rand}(k)$ are the corresponding quantities averaged across 1,000 randomized control networks.

To create randomized control networks, we first rewired the pruned graphs while preserving their degree distribution, and then randomly shuffled the edge weights among the new edges (n = 1,000). Note that the normalization ensures that $\phi_{norm}(k) > 1$ indicates a true weighted rich-club structure, i.e., that highly connected nodes allocate more weight to their mutual connections than expected by chance.

We were mainly interested in whether rich-clubs remained stable over time. Thus, we assessed their existence by aggregating the social networks over consecutive three-day periods: a 1-day resolution made visualization highly impractical, while a 7-day resolution was too coarse to capture meaningful variations. Although the specific members of the rich-club varied across time windows, some individuals were consistently part of it. We defined an animal as part of the 'stable rich-club' (sRC) if it was included in the rich-club in at least 80% of the networks (i.e., in 4 out of 5 graphs). For practical reasons, this analysis was conducted on social interaction networks; however, similar sRCs (~80% overlap) were observed in networks based on approaches and interaction durations (see Supplementary Fig. 7).

**Null model testing.** To assess the significance of our findings on rich-clubs and their temporal stability, we employed null model testing, comparing the experimentally observed values to distributions generated under random conditions to obtain the statistical significance of our results. Each null model represents the expected outcome under the assumption of no systematic relationship between group membership (e.g., OXTR$^{\Delta AON}$ vs. controls, or littermates vs. no littermates…) and rich-club inclusion. The specific implementation of the null model depended on the hypothesis being tested (e.g., mutant exclusion, influence of family ties), but all were based on 10,000 random permutations of relevant experimental labels (e.g., IDs, litter indices, round number).

Exclusion of OXTR$^{\Delta AON}$ mice from the sRC: Specifically, to test whether OXTR$^{\Delta AON}$ mice were excluded from the sRC more often than expected by chance, we randomly reassigned IDs within each group 10,000 times. For each iteration, we counted how many randomized IDs of OXTR$^{\Delta AON}$ mice occupied positions corresponding to sRC members identified experimentally. This null model assumes that all mice within a group have an equal probability of being part of the sRC. Significance was assessed by counting the number of iterations in which the number of randomized OXTR$^{\Delta AON}$ indices in the sRC was less than or equal to the empirical value.

Influence of family ties: To examine whether sRC members were more likely to come from the same litter, we randomized litter indices within each group 10,000 times. For each iteration, we counted the number of groups in which at least two sRC members obtained the same litter index. Statistical significance was assessed by counting how many iterations produced a number of randomized groups with littermates in the sRC that was less than or equal to the number observed in the actual experiment.

Stability of sRC membership across group reassignments: To assess whether sRC membership reflects a stable individual trait or a characteristic of group-level dynamics, we tested whether mice that had previously been part of the sRC were more likely to re-enter it after being reassigned to a different NoSeMaze group. Notably, the group reshuffling procedure ensured that animals were housed with different group members in each NoSeMaze round. We first identified all mice that (1) participated in at least two NoSeMaze rounds, and (2) had been part of the sRC at least once. For each of these mice, we looked at every later round in which they were placed into a new NoSeMaze group. After randomly reassigning IDs within that group, we checked whether the mouse again became part of the sRC. To avoid potential bias in defining the 'initial' sRC membership, one of the eligible rounds was randomly selected as the reference round for each mouse. This null model was repeated across 10,000 independent iterations, each time recording how many of these previously sRC-assigned mice re-entered the sRC at least once following reshuffling. The resulting distribution served as a null model of sRC re-entry, representing the level of membership conservation expected by chance under the assumption that group membership is unrelated to individual traits.

**Graph-theoretical metrics.** To investigate how sRC members differ from $OXTR^{\Delta AON}$ and control animals that were not part of the sRC, we computed three graph-theoretical metrics: out-strength, mean normalized edge fluctuation (NEF), and burstiness. NEF and burstiness [8] were calculated separately for incoming and outgoing directionality using the Python implementation of the 'igraph' toolbox [9] or custom-implemented scripts with Python's NumPy package (see below). The corresponding code is available at https://github.com/KelschLAB/NoSeMaze-stableRichClubs.git. In brief, out-strength captures the total number of approaches initiated by an individual, NEF provides a measure of how strongly an individual's social ties fluctuate over time, and burstiness reflects whether interactions occur at regular intervals or in sudden clusters. Detailed descriptions and formulas for each metric are provided below. All metrics were computed on a daily basis for each individual without aggregating the social networks, as their temporal dynamics provide meaningful insights. To better disentangle the behavior of $OXTR^{\Delta AON}$ animals from the reactions of their social interaction partners, analyses were based on directed graphs constructed from approach networks. Group differences in the graph metrics (sRC members, non-sRC controls, and $OXTR^{\Delta AON}$) were assessed using permutation tests (n = 10,000) based on group medians. To account for repeated measures, group labels were shuffled at the level of the animal, ensuring that all observations from the same individual remained assigned to the same group.

Out-strength: For node $i$, $s_i^{out}$ quantifies the cumulative strength of its outgoing connections, i.e, the total weight of edges from node $i$ to its neighbors:

$$s_i^{out} = \sum_{j \in N_i^{out}} w_{ij}$$

where $N_i^{out}$ is the set of nodes that receive edges from $i$, and $w_{ij}$ is the weight of the edges from $i$ to $j$. We calculated both the mean out-strength across time and the standard deviation of its temporal derivative, with the latter reflecting the variability in an individual's motivation to initiate interactions.

Mean Normalized Edge Fluctuation (NEF): In contrast to the standard deviation of the derivative of strength, which captures fluctuations in an individual's overall activity level, NEF assesses variability at the level of individual connections, thereby providing a more fine-grained measure of temporal instability in social ties. For node $i$ in a network of $N$ nodes, NEF is defined as:

$$NEF_i = \frac{1}{N} \sum_{j \neq i} \frac{\sigma[e_{ij}(t)]}{\mu[e_{ij}(t)]}$$

where $\sigma$ and μ denote the standard deviation and mean, and $e_{ij}(t)$ is the time-series obtained by taking the values of the edge between $i$ and $j$ as a function of time. In other words, NEF corresponds to the average of the coefficient of variation of the temporal evolution of each edge connected to node $i$. It provides a node-level summary of how variable an individual's connections are over time, normalized by the average magnitude of these connections. For directed networks, NEF was computed separately for outgoing edges (from $i$ to others) and incoming edges (from others to $i$).

Mean Burstiness: The mean burstiness can quantify the temporal irregularity of edge activity of a node in a network (Goh & Barabási, 2008). For a binarized graph (non-weighted), the burstiness computation of an edge starts by taking the sequence of inter-contact intervals, i.e. the series of interval duration between the times at which the edge was active. Then, it is obtained via

$$B = \frac{(\sigma - \mu)}{(\sigma + \mu)}$$

where $\sigma$ and μ are the mean and standard deviation of the inter-contact series intervals, respectively. The inter-contact series of an edge between node $i$ and $j$ is given by:

$$\tau_{ij}(k) = t_{ij}(k+1) - t_{ij}(k)$$

where $t_{ij}(1), \dots, t_{ij}(n)$ represent the times at which the edge was active. The resulting value ranges from –1 (perfectly regular) to 1 (highly bursty). The mean burstiness for node $i$ is then obtained by averaging the burstiness between $i$ and all its neighbors. For directed networks, it can be computed distinctly for outgoing edges (from node $i$ to all other nodes) or incoming edges (from all other nodes towards $i$).

**Software and statistical analysis.** Data processing was performed using custom-written scripts in Matlab (Mathworks), R (https://www.r-project.org/, v4.4.1), and Python (https://www.python.org, v3.13). Information about statistical tests used is given either in the related method sections or the figure legends. LME designs are reported in the respective method section and were performed in Matlab. Social network analyses were conducted in Python using the 'igraph' [9] and 'networkX' [10] toolboxes and the statistical analysis of the results was conducted using the Numpy, Pandas, and Scipy packages. Note that all permutation tests accounted for the repeated presence of animals across multiple rounds by shuffling at the level of the animal, ensuring that all observations from the same individual remained assigned to the same group.

**Supplementary References**


1. Fulenwider HD, Caruso MA, Ryabinin AE. Manifestations of domination: Assessments of social dominance in rodents. Genes Brain Behav. 2022;21:e12731.

2. Friard O, Gamba M. BORIS: a free, versatile open-source event-logging software for video/audio coding and live observations. Methods Ecol Evol. 2016;7:1325–1330.

3. Reinwald, JR, Ghanayem, S, Wolf, D, Lebedeva, J, Lebhardt, P, Gölz. O, et al. Individual differences drive social hierarchies in mouse societies. bioRxiv. 2025. 2025.

4. Erskine A, Bus T, Herb JT, Schaefer AT. AutonoMouse: High throughput operant conditioning reveals progressive impairment with graded olfactory bulb lesions. PloS One. 2019;14:e0211571.

5. DAVID HA. Ranking from unbalanced paired-comparison data. Biometrika. 1987;74:432–436.

6. Mathis A, Mamidanna P, Cury KM, Abe T, Murthy VN, Mathis MW, et al. DeepLabCut: markerless pose estimation of user-defined body parts with deep learning. Nat Neurosci. 2018;21:1281–1289.

7. Alstott J, Panzarasa P, Rubinov M, Bullmore ET, Vértes PE. A unifying framework for measuring weighted rich clubs. Sci Rep. 2014;4:7258.

8. Goh K-I, Barabási A-L. Burstiness and memory in complex systems. Europhys Lett. 2008;81:48002.

9. Csárdi G, Nepusz T. The igraph software package for complex network research. InterJournal Complex Syst. 2006:1695.

10. Hagberg A, Swart PJ, Schult DA. Exploring network structure, dynamics, and function using NetworkX. Los Alamos National Laboratory (LANL); 2007.
